# A Sir2-regulated locus control region in the recombination enhancer of *Saccharomyces cerevisiae* specifies chromosome III structure

**DOI:** 10.1101/558494

**Authors:** Mingguang Li, Ryan D. Fine, Manikarna Dinda, Stefan Bekiranov, Jeffrey S. Smith

**Author notes:** Corresponding Author: Department of Biochemistry and Molecular Genetics, University of Virginia School of Medicine, Pinn Hall, Box 800733, Charlottesville, VA 22908. Equally contributed to the work (Mingguang Li, Ryan D. Fine).

## Abstract

The NAD^+^-dependent histone deacetylase Sir2 was originally identified in *Saccharomyces cerevisiae* as a silencing factor for *HML* and *HMR*, the heterochromatic cassettes utilized as donor templates during mating-type switching. *MAT*a cells preferentially switch to *MAT*α using *HML* as the donor, which is driven by an adjacent *cis*-acting element called the recombination enhancer (RE). In this study we demonstrate that Sir2 and the condensin complex are recruited to the RE exclusively in *MAT*a cells, specifically to the promoter of a small gene within the right half of the RE known as *RDT1*. We go on to demonstrate that the *RDT1* promoter functions as a locus control region (LCR) that regulates both transcription and long-range chromatin interactions. Sir2 represses the transcription of *RDT1* until it is redistributed to a dsDNA break at the MAT locus induced by the HO endonuclease during mating-type switching. Condensin is also recruited to the *RDT1* promoter and is displaced upon HO induction, but does not significantly repress *RDT1* transcription. Instead condensin appears to promote mating-type switching efficiency and donor preference by maintaining proper chromosome III architecture, which is defined by the interaction of *HML* with the right arm of chromosome III, including *MAT*a and *HMR*. Remarkably, eliminating Sir2 and condensin recruitment to the *RDT1* promoter disrupts this structure and reveals an aberrant interaction between *MAT*a and *HMR*, consistent with the partially defective donor preference for this mutant. Global condensin subunit depletion also impairs mating type switching efficiency and donor preference, suggesting that modulation of chromosome architecture plays a significant role in controlling mating type switching, thus providing a novel model for dissecting condensin function *in vivo*.

**Author summary:** Sir2 is a highly conserved NAD^+^-dependent protein deacetylase and defining member of the sirtuin protein family. It was identified about 40 years ago in the budding yeast, *Saccharomyces cerevisiae*, as a gene required for silencing of the cryptic mating-type loci, *HML* and *HMR*. These heterochromatic cassettes are utilized as templates for mating-type switching, whereby a programmed DNA double-strand break at the *MAT*a or *MAT*α locus is repaired by gene conversion to the opposite mating type. The preference for switching to the opposite mating type is called donor preference, and in *MAT*a cells, is driven by a cis-acting DNA element called the recombination enhancer (RE). It was believed that the only role for Sir2 in mating-type switching was silencing HML and HMR. However, in this study we show that Sir2 also regulates expression of a small gene (RDT1) in the RE that is activated during mating-type switching. The promoter of this gene is also bound by the condensin complex, and deleting this region of the RE drastically changes chromosome III structure and alters donor preference. The RE therefore appears to function as a complex locus control region (LCR) that links transcriptional control to chromatin architecture, and thus provides a new model for investigating the underlying mechanistic principles of programmed chromosome architectural dynamics.

## Introduction

Since the first descriptions of mating-type switching in budding yeast approximately 40 years ago, characterization of this process has led to numerous advances in understanding mechanisms of gene silencing (heterochromatin), cell-fate determination (mating-type), and homologous recombination (reviewed in [1]. For example, the NAD^+^-dependent histone deacetylase, Sir2, and other <u>S</u>ilent <u>I</u>nformation <u>R</u>egulator (SIR) proteins, were genetically identified due to their roles in silencing the heterochromatic *HML* and *HMR* loci, which are maintained as silenced copies of the active *MAT*α and *MAT*a loci, respectively [2-4]. The SIR silencing complex (Sir2-Sir3-Sir4) is recruited to cis-acting E and I silencer elements flanking *HML* and *HMR* through physical interactions with silencer binding factors Rap1, ORC, and Abf1, as well as histones H3 and H4 (reviewed in [5]).

*HML* and *HMR* play a critical role in mating-type switching. Haploid cells of the same mating-type cannot mate to form diploids, the preferred cell type in the wild. Therefore, in order to facilitate mating and diploid formation, haploid mother cells switch their mating type by expressing HO endonuclease, which introduces a programmed DNA double-strand break (DSB) at the *MAT* locus [6]. The break is then repaired by homologous recombination using either *HML* or *HMR* as a donor template for gene conversion [6, 7]. This change in mating type enables immediate diploid formation between mother and daughter. *HO* is deleted from most standard lab strains in order to maintain them as haploids, so expression of *HO* from an inducible promoter such as P_*GALI*_ is commonly used to switch mating types during strain construction [8].

There is a “donor preference” directionality to mating-type switching such that ∼90% of the time, the HO-induced DSB is repaired to the opposite mating type [9]. For example, *MAT*α cells preferentially switch to *MAT*a using *HMR* as the donor. However, while both silent mating loci can be utilized as a donor template, usage of *HML* by *MAT*a cells requires a 2.5 kb intergenic region located ∼17 kb from *HML* called the recombination enhancer (RE) [10]. Donor preference activity within the RE has been further narrowed down to a 700 bp segment containing an Mcm1/α2 binding site (DPS1) and multiple Fkh1 binding sites [10]. The RE is active in *MAT*a cells, requiring Mcm1 and Fkh1 activity at their respective binding sites [10-12]. The RE is inactivated in *MAT*α cells due to expression of transcription factor α2 from *MAT*α [13], which forms a repressive heterodimer with Mcm1 (Mcm1/α2) to repress *MAT*a-specific genes [1]. Current models for donor preference posit that Fkh1 at the RE helps position *HML* in close proximity with *MAT*a by interacting with threonine-phosphorylated H2A (γ-H2AX) and Mph1 DNA helicase at the HO-induced DNA DSB [14, 15].

Sir2-dependent silencing of *HML* and *HMR* has two known functions related to mating-type switching. First, *HML* and *HMR* must be silenced in haploids to prevent formation of the a1/α2 heterodimer, which would otherwise inactivate haploid-specific genes such as *HO* [16]. Second, heterochromatin structure at *HML* and *HMR* blocks cleavage by HO, thus restricting its activity to the fully accessible *MAT* locus [17, 18]. Here we describe new roles for Sir2 and the condensin complex within the RE during mating-type switching. ChIP-seq analysis revealed strong overlapping binding sites for Sir2 and condensin at the promoter of a small gene within the RE known as *RDT1*. Sir2 was found to repress the *MAT*a-specific transcription of *RDT1*, which is also translated into a small 28 amino acid peptide. *RDT1* expression is also dramatically upregulated during mating-type switching when Sir2 redistributes to the HO-induced DNA DSB at *MAT*a. Furthermore, eliminating Sir2/condensin recruitment to the *RDT1* promoter disrupts chromosome III architecture such that mating-type switching efficiency and donor preference are partially impaired. The *RDT1* promoter region therefore functions like a classic locus control region (LCR) in *MAT*a yeast cells, regulating localized transcription as well as long-range chromosome interactions.

## Results

### Sir2 and condensin associate with the recombination enhancer (RE)

We previously characterized global sirtuin distribution using ChIP-Seq to identify novel loci regulated by Sir2 and its homologs [19]. Significant overlap was observed between binding sites for Sir2, Hst1, or Sum1 with previously described condensin binding sites [19, 20], suggesting a possible functional connection. ChIP-Seq was therefore performed on WT and *sir2*Δ strains in which the condensin subunit Smc4 was C-terminally tagged (13xMyc) (Fig 1A). To avoid “hyper-ChIPable” loci that can appear in yeast ChIP-seq experiments, we also ran nuclear localized GFP controls [21]. Genes closest to Sir2-dependent condensin peaks after subtraction of GFP are listed in Table S1, and are distributed throughout the genome. One of the strongest peaks overlapped with a Sir2-myc binding site on chromosome III between *KAR4* and *SPB1* that was not enriched for GFP (Fig 1A). The specificity of Sir2 enrichment at this peak, as opposed to the adjacent *SPB1* gene, was independently confirmed by quantitative ChIP using an α-Sir2 antibody (Fig 1B), with enrichment comparable to levels observed at the *HML-I* silencer (Fig 1A and B). Sir2-dependent condensin binding was also confirmed for Myc-tagged Smc4 and Brn1 subunits (Fig 1C). The ∼2.5 kb intergenic region between *KAR4* and *SPB1* was previously defined as a cis-acting recombination enhancer (RE) that specifies donor preference of mating-type switching in *MAT*a cells [10, 13]. Quantitative ChIP assays revealed that Sir2 and Brn1-myc enrichment at the RE was also *MAT*a-specific (Fig 1D and E), which was notable because the ChIP-seq datasets in Fig 1A happened to be generated from *MAT*a strains. We next considered whether condensin binding in the *MAT*a *sir2*Δ mutant was due to *HMLALPHA2* expression caused by defective *HML* silencing. To test this idea, we retested Brn1-myc ChIP at the RE in strains lacking *HML*, and found that deleting *SIR2* no longer affected condensin recruitment (Fig 1F). Similarly, a *MAT*a condensin mutant (*ycs4-1*) known to have an *HML* silencing defect [22] reduced Sir2 recruitment to the RE, but had no effect when *HML* was deleted (Fig 1G). Sir2 and condensin are therefore independently recruited to the RE specifically in *MAT*a cells.

**Fig 1.**
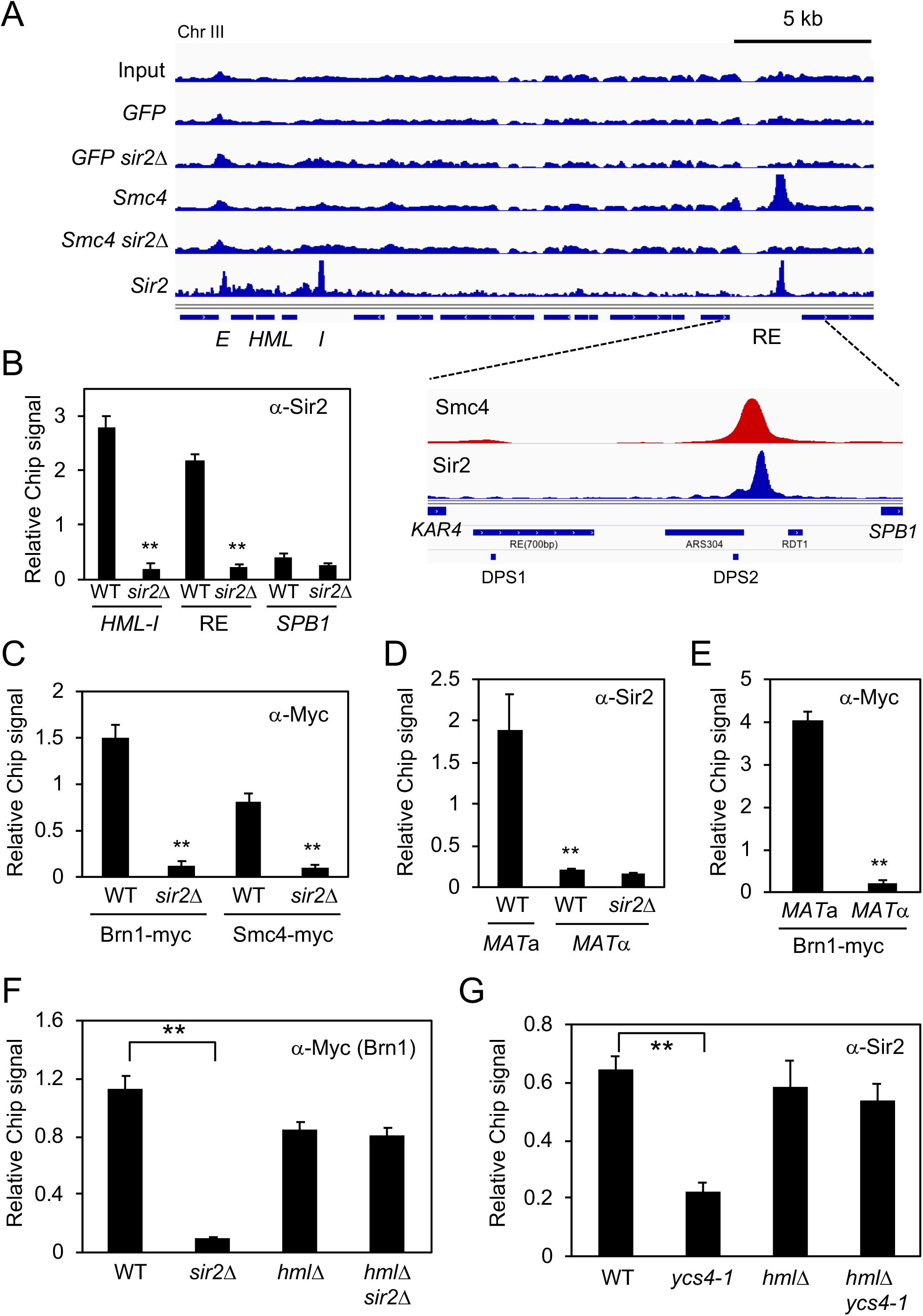
*MAT*a-specific binding of Sir2 and condensin to the recombination enhancer (RE). **(A)** Chip-seq of Smc4-myc, Sir2-myc, and nuclear localized GFP in WT and *sir2*Δ backgrounds. The left arm of chromosome III is depicted from *HML* to *SPB1*. RE indicates the recombination enhancer region. Inset: The minimal 700bp RE element required for donor preference is indicated, as are the two Mcm1/α2 binding sites (DPS1 and DPS2) and *RDT1*. **(B)** Sir2 ChIP at the RE, *HML-I* silencer, and *SPB1*. **(C)** α-Myc ChIP of Brn1-myc and Smc4-myc at the RE. **(D)** ChIP showing *MAT*a-specific binding of Sir2 to the RE. **(E)** ChIP showing *MAT*a-specific binding of Brn1-myc to the RE. **(F)** Brn1-Myc ChIP at RE is not Sir2 dependent. **(G)** Native Sir2 ChIP at RE is not condensin dependent. ChIP signal relative to input is plotted as the mean of three replicates. Error bars = standard deviation. (**p<0.005).

### Sir2 regulates a small gene (*RDT1*) within the RE

Donor preference activity ascribed to the RE was previously narrowed down to a *KAR4 (YCL055W)*-proximal 700 bp domain defined by an Mcm1/α2 binding site (Fig 2A, *DPS1*) [10, 11, 13]. The Sir2 and condensin ChIP-seq peaks we identified were located outside this region, between a second Mcm1/α2 binding site (*DPS2*) and a small gene of unknown function called *RDT1* [23] (Fig 1A and 2A). We noticed the location of *RDT1* coincided with the smallest of several putative non-coding RNAs (ncRNA) previously reported as being transcribed from the RE, but not annotated in SGD [24, Fig 2A]. Quantitative RT-PCR and analysis of publicly available RNA-seq data from BY4741 (*MAT*a) and BY4742 (*MAT*α) revealed that *RDT1* expression was indeed *MAT*a specific (Fig 2B and S1A).

**Fig 2.**
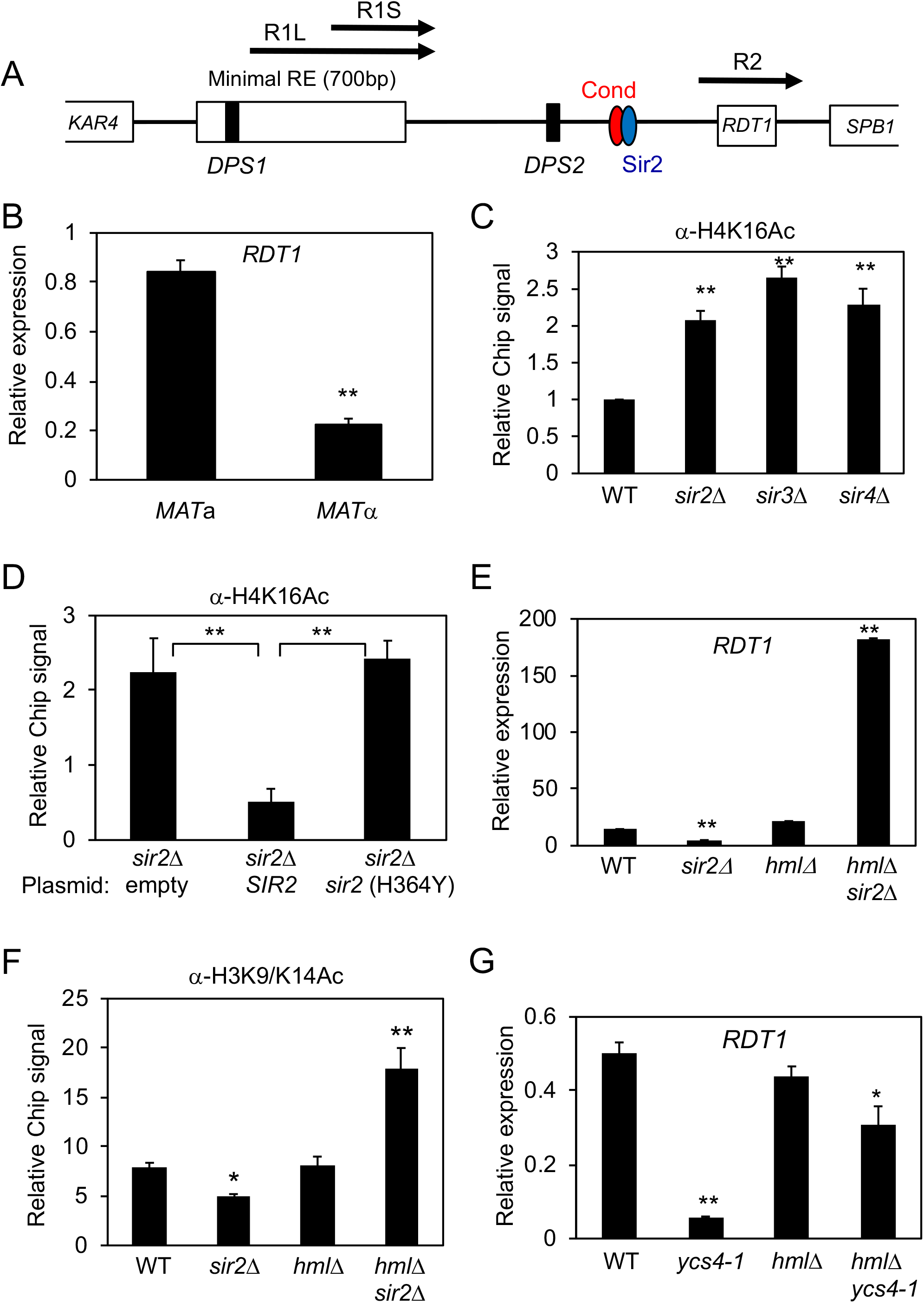
*RDT1* is a novel Sir2 regulated gene. **(A)** Schematic of RE locus depicting Sir2/Condensin peak location relative to previously reported R1L/S and R2 RNA (*RDT1*). **(B)** *RDT1* mRNA expression is *MAT*a specific. **(C)** H4K16ac ChIP at RE in SIR complex null strains. **(D)** H4K16ac deacetylation is dependent on Sir2 catalytic activity. A *sir2*Δ strain was transformed with the indicated plasmids and ChIP assays performed. **(E)** Differential *RDT1* transcriptional regulation by *SIR2* is dependent on *HML* status. **(F)** Effect of *sir2*Δ on H3K9/K14ac ChIP at the *RDT1* promoter in *HML* and *hml*Δ backgrounds. **(G)** Effects of the temperature sensitive *ycs4-1* mutation on *RDT1* expression in *HML* and *hml*Δ backgrounds. (*p<0.05; **p<0.005).

We next asked whether Sir2 and/or condensin regulate histone acetylation and *RDT1* expression when recruited to the RE. Sir2 normally represses transcription at *HML, HMR*, and telomeres as a catalytic subunit of the SIR complex where it preferentially deacetylates H4K16 (reviewed in [5]). Accordingly, deleting *SIR2, SIR3*, or *SIR4* from *MAT*a cells increased H4K16 acetylation at the *RDT1* promoter (Fig 2C), consistent with the observed enrichment of Sir3-myc and Sir4-myc at this site (Fig S1B). Furthermore, re-introducing active *SIR2* into the *sir2*Δ mutant restored H4K16 to the hypoacetylated state, whereas catalytically inactive *sir2-H364Y* did not (Fig 2D).

Deleting *SIR2* initially appeared to repress *RDT1* expression in *MAT*a cells (Fig 2E), but we hypothesized this was due to *HMLALPHA2* derepression and formation of the Mcm1/α2 repressor, which could locally repress *RDT1* through the adjacent Mcm1/α2 binding sites. Indeed, simultaneously deleting *SIR2* and *HML* resulted in very high *RDT1* expression (Fig 2E), which was increased even further when the paralogous *HST1* gene was also deleted (Fig S1C), indicating some redundancy. By eliminating *HML* we also observed elevated histone H3 acetylation in the absence of *SIR2* (Fig 2F), providing strong evidence that the SIR complex establishes a generally hypoacetylated chromatin environment at the *RDT1* promoter that requires effective silencing at *HML*. On the other hand, *RDT1* was not upregulated in an *hml*Δ *ycs4-1* condensin mutant (Fig 2G), suggesting that condensin has a different functional role at this locus.

We next attempted to block Sir2 and condensin recruitment to the *RDT1* promoter by precisely deleting a 100bp DNA sequence underlying the shared enrichment region (coordinates 30701-30800), while not disturbing the adjacent Mcm1/α2 site (Fig 3A). Sir2 and Brn1-myc binding to the RE as measured by ChIP was greatly diminished in this mutant (Fig 3B and 3C), despite unaltered Sir2, Brn1-myc, or Smc4-myc expression levels (Fig S2A-C). Furthermore, *RDT1* transcriptional expression was significantly increased by the 100bp deletion exclusively in *MAT*a cells (Fig 3D), consistent with the loss of Sir2-mediated repression.

**Fig 3.**
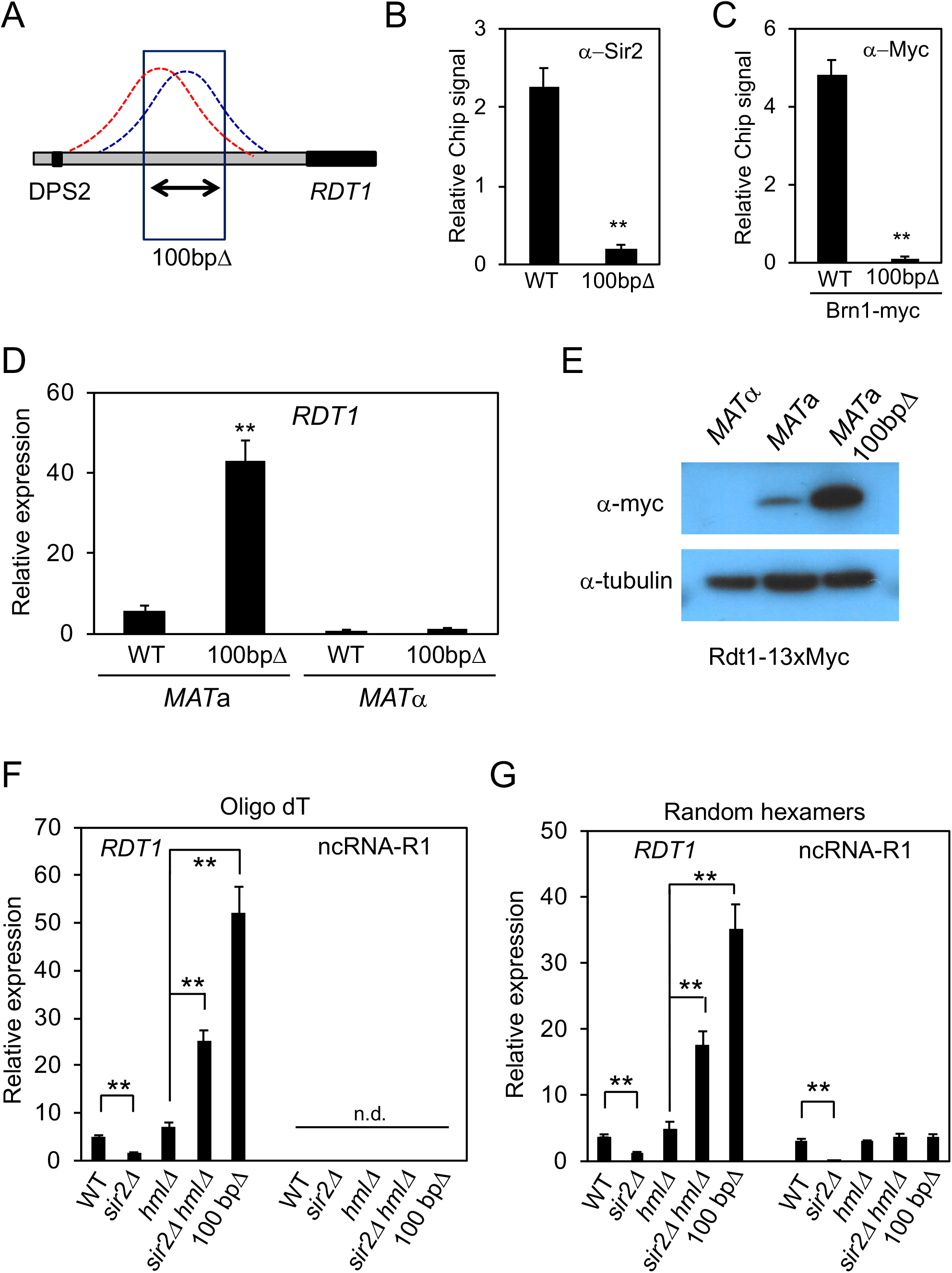
Identification of a 100bp sequence that recruits Sir2/condensin and represses RDT1 expression. **(A)** Schematic indicating a 100bp deletion that covers the condensin (red) and Sir2 (blue) peaks. (B) ChIP of Sir2 in the 100bpΔ mutant (ML275). **(C)** ChIP of Brn1-Myc in the 100bpΔ mutant. **(D)** *RDT1* transcription in *MAT*a cells is derepressed in the 100bpΔ mutant. **(E)** Western blot of Rdt1-13xMyc in WT *MAT*α and *MAT*a cells, as well as the *MAT*a 100bpΔ mutant. **(F)** *RDT1* and R1 expression when using oligo dT priming for the reverse transcription step. **(G)** *RDT1* and R1 expression when using random hexamer priming for reverse transcription. (**p<0.005).

Because Sir2 and condensin were not present at the *RDT1* promoter in *MAT*α cells, we reasoned that their binding should require a *MAT*a specific transcription factor. This made the 2^nd^ Mcm1/α2 binding site (DPS2) upstream of the Sir2/condensin ChIP-seq peaks an ideal candidate because it has not been ascribed a function other than redundancy with DPS1. Deleting *MCM1* is lethal, so alternatively, we deleted the 2^nd^ Mcm1/α2 binding site (ChrIII coordinates 30595 to 30626, Fig S3A) and then retested for Sir2 and Brn1-myc enrichment. As shown in Fig S3B and S3C, respectively, Sir2 and Brn1-myc enrichment at both the Mcm1/α2 binding site (DPS2) and the *RDT1* promoter (defined as the Sir2/condensin peaks) was significantly reduced in the binding site mutant. These results suggest that Mcm1 may nucleate a complex that recruits the SIR and condensin complexes to the *RDT1* promoter in *MAT*a cells, and also provides a possible mechanism of blocking the recruitment in *MAT*α cells due to the interaction of Mcm1 with α2.

### *RDT1* encodes a translated mRNA

<u>R</u>ibosome <u>D</u>etected <u>T</u>ranscript-1 (*RDT1*) was originally annotated as a newly evolved gene whose transcript was associated with ribosomes and predicted to have a small open reading frame of 28 amino acids [23]. Our work suggested that *RDT1* and the putative non-coding R2 transcript were the same (Fig 2A). To determine if *RDT1*/R2 codes for a small protein, the ORF was C-terminally fused with a 13x-Myc epitope in *MAT*a and *MAT*α cells. As shown in Fig 3E, a fusion protein was detectable in exponentially growing *MAT*a WT cells and also highly expressed in the 100bpΔ background, correlating with the increased RNA level observed for that mutant in Fig 3D.

Additional *MAT*a-specific RNAs are derived from the minimal 700bp RE domain (Fig 2A; R1L and R1S) [13, 25], so we next tested whether Sir2 controls their expression from a distance. As shown in Fig 3F, qRT-PCR using standard oligo(dT) primers for cDNA synthesis effectively measured *RDT1* expression at predicted levels for the various strains tested, but the R1 RNAs were not detectable. Many long non-coding RNAs (lncRNAs) are not polyadenylated [26], so the cDNA synthesis was repeated using random hexamer primers. In *MAT*a WT cells (ML1), R1L/S RNAs were now detected at levels comparable to *RDT1* (Fig 3G). Similar to *RDT1*, R1L/S RNAs were repressed in the absence of *SIR2* due to the *HMLALPHA2* pseudodiploid derepression phenotype. But unlike *RDT1*, the R1L/S RNA expression level was not elevated in the 100bpΔ or *hml*Δ *sir2*Δ mutants, indicating these RNAs are not under direct Sir2 control, but are strongly repressed in the absence of Sir2. We conclude that the R1L/S RNAs are most likely non-polyadenylated lncRNAs, whereas *RDT1* is Sir2-repressed and polyadenylated mRNA that can be translated into a small protein of unknown function.

### Sir2 and condensin are displaced from the *RDT1* promoter during mating-type switching

We next asked if Sir2 played any role in regulating *RDT1* during mating-type switching. Sir2 was previously shown to associate with a HO-induced DSB at the *MAT* locus during mating-type switching, presumably to effect repair through histone deacetylation [27]. Transient Sir2 recruitment to the DSB could potentially occur at expense of the *RDT1* promoter, thus resulting in *RDT1* derepression. To test this idea, HO was induced at time 0 with galactose and then turned off 2 hours later by glucose addition to allow for repair/switching to occur (Fig 4A and B). By the 3 hr time point (1hr after glucose addition), ChIP analysis indicated Sir2 was maximally enriched at the *MAT* locus (Fig 4C), corresponding to the time of peak mating-type switching ([27] and Fig 4B). Interestingly. Sir2 was significantly depleted from the *RDT1* promoter within 1 hr after HO induction, and by 3 hr there was actually stronger enrichment of Sir2 at *MAT* than *RDT1* (Fig 4C). Critically, this apparent Sir2 redistribution coincided with maximal induction of *RDT1* mRNA and the Myc-tagged Rdt1protein (Fig 4D and 4E, 3 hr). Once switching was completed by 4 hr (2hr after glucose addition), *RDT1* transcription was permanently inactivated and Sir2 binding never returned because most cells were now *MAT*α. The Myc-tagged Rdt1 protein, however, remained elevated for the rest of the time course (Fig 4E), suggesting that it is relatively stable, at least when epitope tagged. A parallel ChIP time course experiment was performed with condensin (Brn1-myc), resulting in significant depletion from the *RDT1* promoter within 1 hr (Fig 4F), similar to the timing of Sir2 loss. However, rather than redistributing to the DSB, Brn1-myc enrichment was actually reduced at the break site, suggesting that condensin normally associates with *MAT*a in non-switching cells, but becomes displaced in response to the HO-induced DSB, perhaps to facilitate structural reorganization associated with switching.

**Fig 4.**
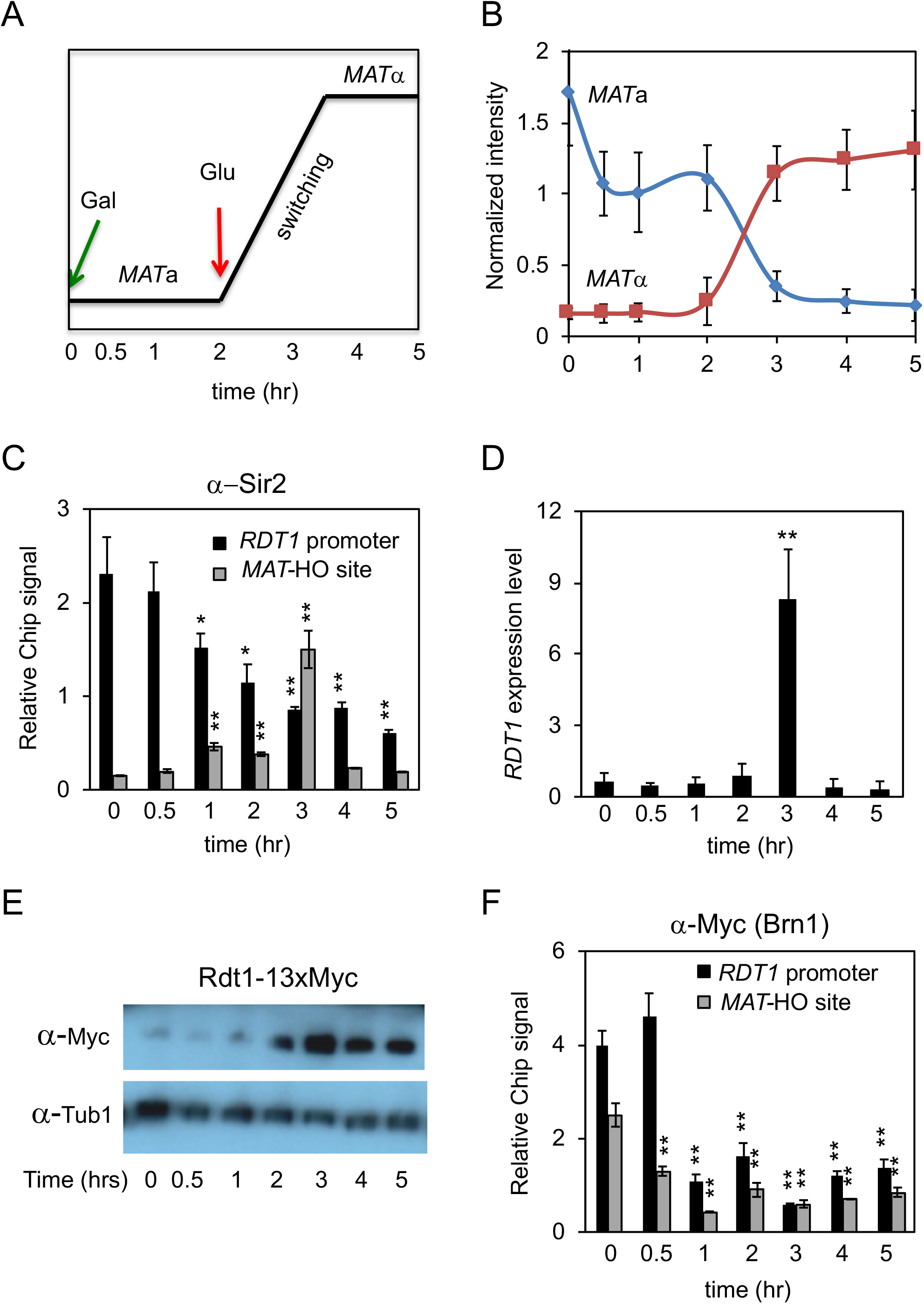
Dynamics of Sir2 and condensin binding at the *RDT1* promoter and *MAT*a locus during mating-type switching. **(A, B)** Mating-type switching time course where HO was induced by galactose at time 0, then glucose added at 2 hr to stop HO expression and allow for break repair. Switching is maximal at 3 hr [27]. **(C)** ChIP of Sir2 at the *RDT1* promoter and the HO-induced DSB (*MAT*-HO). **(D)** qRT-PCR of *RDT1* expression across the mating-type switching time course. **(E)** Rdt1-13xMyc protein expression across the same time course. **(F)** ChIP of Brn1-myc at the *RDT1* promoter and MAT-HO break site across the same time course. (*p<0.05, **p<0.005 compared to time 0).

### The *RDT1* promoter region controls chromosome III architecture

The coupling of Sir2 and condensin distribution with *RDT1* transcriptional regulation during mating-type switching was reminiscent of classic locus control regions (LCR) that modulate long-range chromatin interactions. We therefore hypothesized that the *RDT1* promoter region may function as an LCR to modulate long-range chromatin interactions of chromosome III. To test this hypothesis, we performed Hi-C analysis with WT, *sir2*Δ and the 100bpΔ strains. Genomic contact differences between the mutants and WT were quantified using the HOMER Hi-C software suite [28], and the frequency of statistically significant differences for each chromosome calculated (Fig 5A). Chromosome III had the most significant differences in both mutants, so we focused on this chromosome and used HOMER to plot the observed/expected interaction frequency in 10kb bins for each strain as a heat map (Fig 5B). In a WT strain (ML1) there was strong interaction between the left and right ends of chromosome III, mostly centered around the *HML* (bin 2) and *HMR* (bin 29) loci. Interestingly, *HML* (bin 2) also appeared to sample the entire right arm of chromosome III, with the interaction frequency increasing as a gradient from *CEN3* to a maximal observed interaction at *HMR*, thus also encompassing the *MAT*a locus at bin 20. This distinct interaction pattern was completely disrupted in the *sir2*Δ mutant, whereas some telomere-telomere contact was retained in the 100bpΔ mutant (Fig 5B), suggesting there was still limited interaction between the left and right ends of the chromosome. We confirmed the changes in *HML*-*HMR* interaction for these strains using a quantitative 3C-PCR assay to rule out sequencing artifacts, and also confirmed an earlier *sir2*Δ 3C result from the Dekker lab [29]. Importantly, despite the loss of *HML*-*HMR* interaction in the 100bpΔ mutant, heterochromatin at these domains was unaffected based on normal quantitative mating assays (Fig S4A), and unaltered Sir2 association with *HML* (Fig S4B).

**Fig 5.**
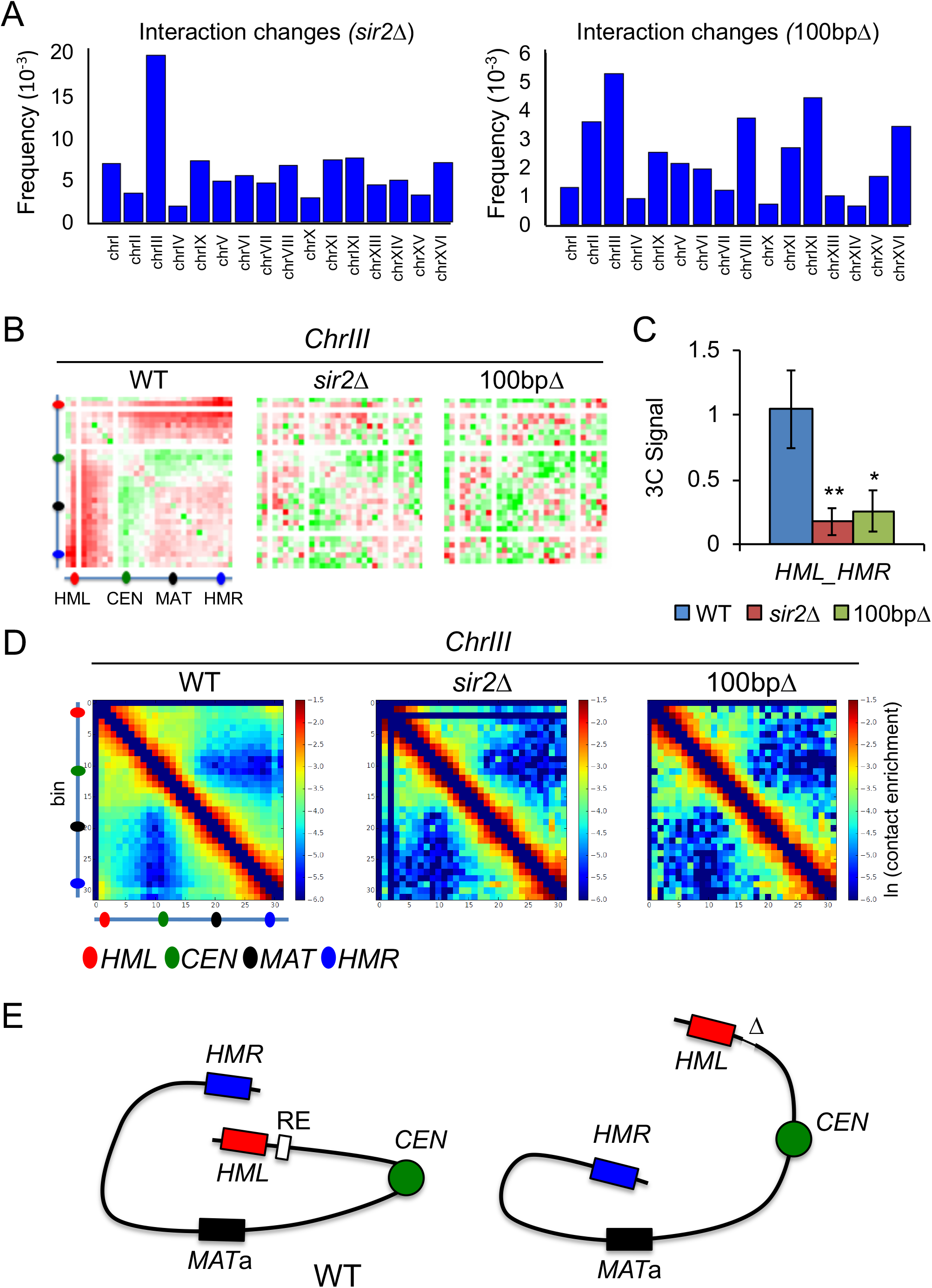
The Sir2/condensin binding site controls chromosome III architecture. Frequency of significant Hi-C interaction changes identified using HOMER for each chromosome in the *sir2*Δ (ML25) and 100bpΔ (ML275) strains compared to WT (ML1). **(B)** HOMER-generated observed/expected Hi-C interaction frequency heat maps (10kb bins) for chromosome III. **(C)** qPCR detection of *HML-HMR* interaction using 3C analysis. (*p < 0.05, **<0.005). **(D)** Iteratively corrected and read-normalized Hi-C heat maps revealing an interaction between *HMR* (bin 29) and *MAT*a (bin 20) in the *sir2*Δ and 100bpΔ mutants. **(E)** Summary of large-scale changes in chromosome III architecture. Δ indicates the 100bp deletion.

We next analyzed the Hi-C data using an iterative correction method that reduces background to reveal interacting loci that potentially drive the overall chromosomal architecture, rather than passenger locus effects [30]. *HML* (bin 2) and *HMR* (bin 29) again formed the dominant interaction pair off the diagonal in WT, which was lost in the *sir2*Δ or 100bpΔ mutants (Fig 5D). Importantly, a prominent new interaction between *HMR* (bin 29) and *MAT*a (bin 20) appeared in both mutants (Fig 5D and E), as would be predicted if normal donor preference of *MAT*a cells was altered. We conclude that the *RDT1* promoter does function like an LCR in *MAT*a yeast cells, regulating localized transcription as well as long-range chromatin interactions relevant to mating-type switching (Fig 5E).

### Sir2 and condensin regulate mating-type switching

Sir2/condensin binding was observed in the right half of the RE (Fig 1A), but this region was previously reported as being dispensable for donor preference activity [10]. Considering that *HMR* was aberrantly associated with the *MAT*a locus in *sir2*Δ and 100bpΔ mutants (Fig 5), we proceeded to test whether these mutants had any alterations in donor preference. A reporter strain was used in which *HMR*a on the right arm of chromosome III was replaced with an *HMR*α allele containing a *Bam*HI site (*HMR*α*-B*) (Fig 6A). After inducing switching to *MAT*α following HO induction with galactose, the proportion of *HML*α or *HMR*α–B utilization for switching was determined by *Bam*HI digestion of a *MAT*α-specific PCR product (Fig 6B) [15]. As expected for normal donor preference, *HMR*α–B on the right arm was only utilized ∼9% of the time in the WT strain, as compared to 91% for *HML*α (Fig 6C). Strikingly, donor preference was completely lost in the *sir2*Δ mutant, similar to a control strain with the RE deleted (Fig 6C and D), and consistent with the clear interaction between *HMR* and *MAT*a observed for the *sir2*Δ mutant in Fig 5D and E. This interaction was less prominent in the 100bpΔ mutant (Fig 5D), and the change in donor preference was also less severe (∼25% *HMR*α-B), though still significantly different from WT (Fig 6C and D). Additionally, we measured the efficiency of switching to *MAT*α across a time course in the ML1 strain background used for ChIP and Hi-C analyses, and did not observe a significant difference between WT and the 100bpΔ mutant. However, switching to *MAT*α was severely impaired in the *sir2*Δ mutant (Fig 6E). We suspect the larger effect on switching efficiency and donor preference in *sir2*Δ cells is due to the derepression of *HMLALPHA2*, because α2 protein normally inactivates the RE in *MAT*α cells [13]. Silencing of *HML* is therefore critical for donor preference in the mating-type switching of *MAT*a cells by preventing expression of the repressive α2 transcription factor.

**Fig 6.**
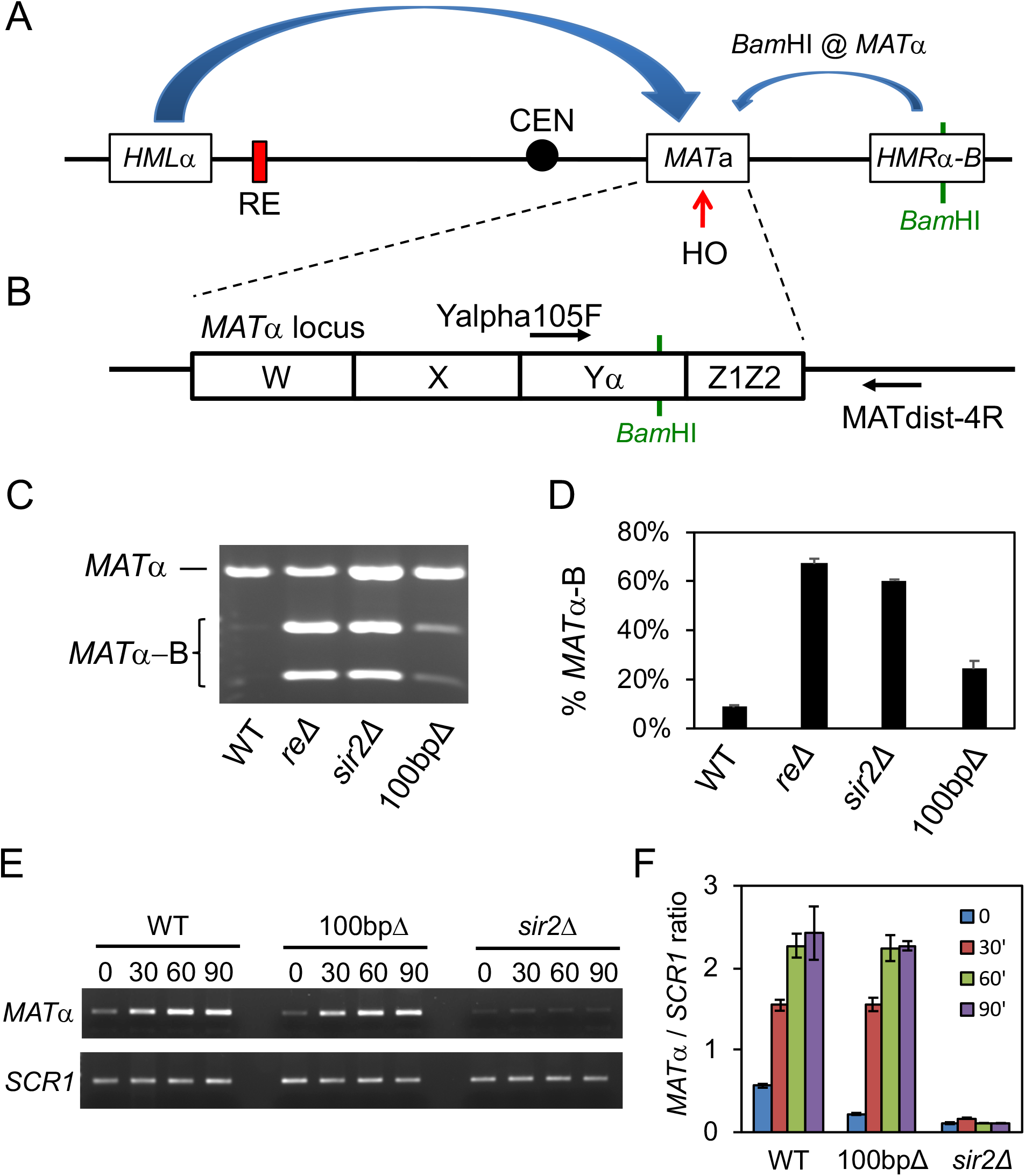
Loss of Sir2 and the Sir2/condensin binding site alters mating-type switching. **(A)** Schematic of a donor preference assay in which utilization of an artificial *HMR*α-B cassette as the donor for switching introduces a unique *Bam*HI site to the *MAT* locus. **(B)** Locations of primers flanking the *Bam*HI site used for PCR detection of *MAT*α. **(C)** Representative ethidium bromide stained agarose gel of *Bam*HI-digested *MAT*α PCR products after mating-type switching in WT (XW652), *re*Δ (XW676), *sir2*Δ (ML557), and 100bpΔ (SY742) strains. The *MAT*α-B product is digested into 2 smaller bands. **(D)** Quantifying the percentage of *MAT*α PCR product digested by *Bam*HI, from three biological replicates. ImageJ was used for the quantitation. (**p < 0.005). **(E)** Time course of switching from *MAT*a to *MAT*α in WT (ML447), 100bpΔ (ML460), and *sir2*Δ (ML458) strains after HO was induced for 45 min and then shut down with glucose. Aliquots were harvested at 30 min intervals. *SCR1* is a control for input genomic DNA. **(F)** ImageJ quantification of *MAT*α PCR relative to *SCR1* for each time point.

Since condensin is also recruited to the *RDT1* promoter region, we were next interested in whether condensin activity was important for mating-type switching. Each gene for the condensin subunits is essential, so instead of using deletions, in the ML1 strain background we C-terminally tagged the Brn1 subunit with an auxin-inducible degron (AID) fused with a V5 epitope. This system allows for rapid depletion of tagged proteins upon addition of auxin to the growth media [31]. Indeed, Brn1-AID was effectively degraded within 15 min of adding auxin (Fig S5A). Importantly, even after 1 hr of auxin treatment, there were no changes in *RDT1* or *HMLALPHA2* gene expression indicated by qRT-PCR (Fig S5B and C), indicating that silencing of *HML* was unaffected, unlike the *ycs4-1* condensin mutant used in Fig 1G [22]. The efficiency of ML1 switching from *MAT*a to *MAT*α was then tested across a time course with or without auxin treatment (Fig 7A). As shown in Fig 7B and C, auxin treatment significantly slowed the pace of switching to *MAT*α, which also suggested there could be a modest effect on donor preference similar to that observed with the 100bpΔ strain. Indeed, Brn1-AID depletion produced a minor, yet significant, alteration in donor preference using the *HMR*α-B reporter strain (Fig 7D). Taken together, these results support a model whereby condensin recruited to the *RDT1* promoter in *MAT*a cells organizes chromosome III into a conformation that favors association of the *MAT*a locus with *HML* instead of *HMR*, thus partially contributing to donor preference regulation.

**Fig 7.**
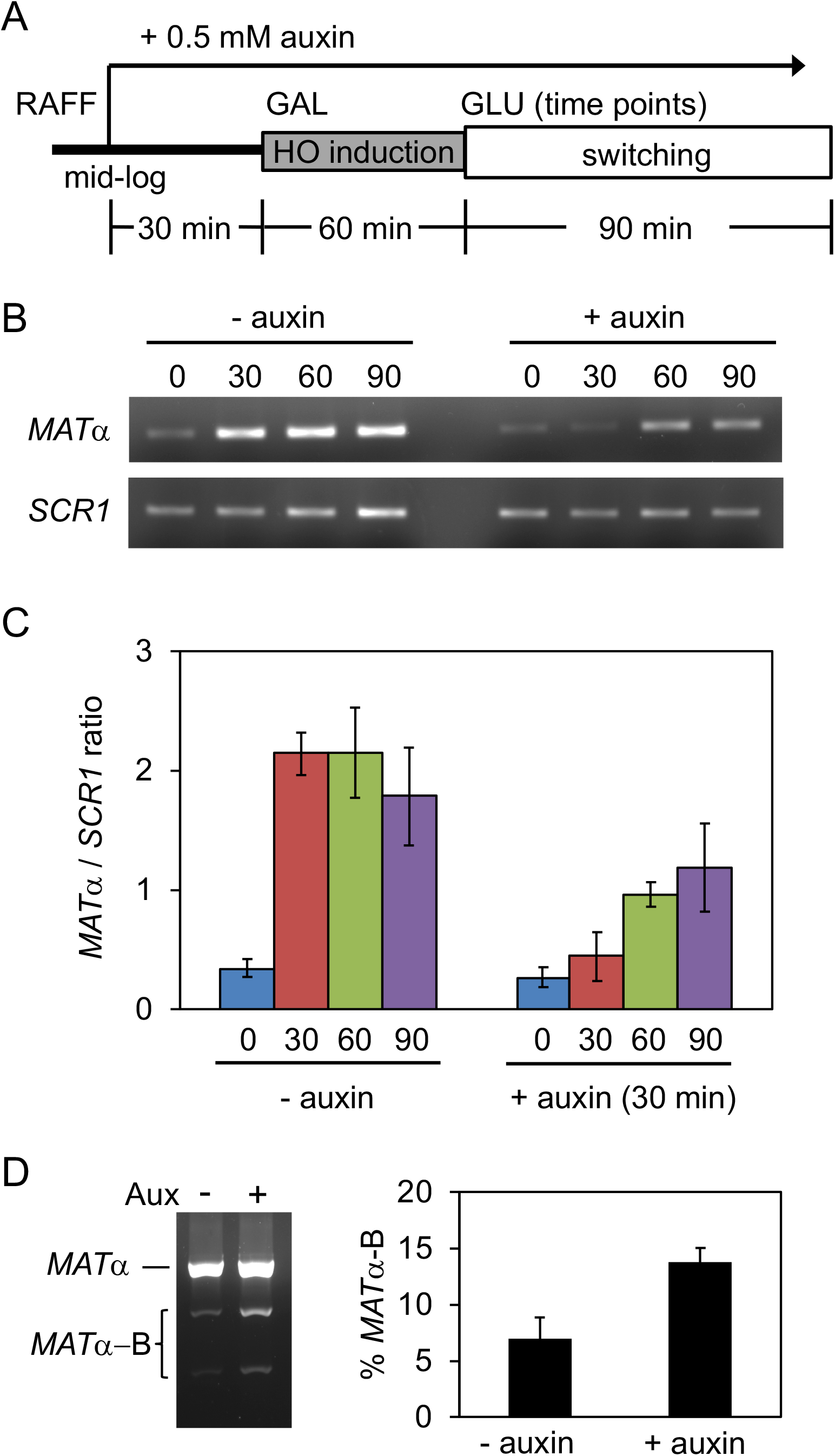
Effects of condensin depletion on mating-type switching. **(A)** Schematic of the time course used to deplete Brn1-AID prior to the induction of mating-type switching in the ML1 strain background. Auxin was added 30 min prior to the induction of HO expression by galactose. **(B)** EtBr stained agarose gel of *MAT*α PCR products amplified from each time point during mating-type switching. *SCR1* PCR was used as a control for input DNA. **(C)** Quantification of the *MAT*α/*SCR1* PCR product ratio across the time course from 3 biological replicates. **(D)** Effect of Brn1-AID depletion on mating-type switching donor preference. A representative biological replicate is shown, along with quantitation of switching using the *HMR*α-B cassette.

## Discussion

*SIR2* was identified almost 40 years ago as a recessive mutation unlinked from *HML* and *HMR* that caused their derepression [3, 4], and has been extensively studied ever since as encoding a heterochromatin factor that functions not only at the *HM* loci, but also telomeres and the rDNA locus (reviewed in [5]). In this study we describe a previously unidentified Sir2 binding site that overlaps with a major non-rDNA condensin binding site within the RE on chromosome III in *MAT*a cells. Here, Sir2 regulates a small gene of unknown function called *RDT1*, which is transcriptionally activated during mating-type switching due to redistribution of repressive Sir2 from the *RDT1* promoter to the HO-induced DSB at *MAT*a. The *RDT1* RNA transcript is also polyadenylated and translated into a small protein, but we have not yet been able to assign a function to the gene or protein because deleting the 28 amino acid ORF had no measurable effect on mating-type switching when using the *GAL-HO* based assays tested thus far (data not shown). It remains possible that deleting *RDT1* would have a significant effect on switching in the context of native HO expression, which is expressed only in mother cells during late G1, whereas the *GAL1-HO* is overexpressed in all cells throughout the cell cycle. It is also possible that *RDT1* functions as a non-coding RNA that happens to be translated into a small non-functional peptide. Alternatively, transcription of *RDT1* could directly function in chromosome III conformation by altering local chromatin accessibility at the promoter. Such a model was proposed for regulation of donor preference by transcription of the R1S/R1L non-coding RNAs [13, 25]. Dissecting the function(s) of *RDT1* therefore remains an area of active investigation for the lab, and perhaps the key to fully understanding how its promoter functions as an LCR.

### Functional complexity within the RE

While we do not yet know the molecular function of *RDT1* in mating-type regulation or other cellular processes, the promoter region of this gene clearly controls the structure of chromosome III. Three-dimensional chromatin structure has long been proposed to influence donor preference [32, 33]. However, deleting the minimal 700bp (left half) of the RE alters donor preference without a large change in chromosome III conformation. Furthermore, deleting the right half of the RE, which includes *RDT1*, changes chromosome III conformation without a dramatic change in donor preference [10, 13, 34]. Based on these findings it was proposed that the RE is a bipartite regulatory element [34], with the left half primarily responsible for donor preference activity and the right half for chromosome III structure. Our results support this view and narrow down the structural regulatory domain of the RE to a small (100bp) region of the *RDT1* promoter bound by the SIR and condensin complexes. Importantly, deleting this small region not only altered chromosome III structure, but also had a significant effect on donor preference, though not as strong as the *sir2*Δ mutation.

The coordination of *RDT1* expression with loss of Sir2/condensin binding at its promoter during mating-type switching, together with the loss of *HML-HMR* interaction in the 100bpΔ mutant, makes this site intriguingly similar to classic locus control regions (LCRs) in metazoans, which are cis-acting domains that contain a mixture of enhancers, insulators, chromatin opening elements, and tissue-specificity elements [35]. The minimal RE was previously described as an LCR in the context of donor preference [10], and transcription of the R1S/R1L long non-coding RNAs via activation by the 1^st^ Mcm1/α2 binding site (DPS1) appears to be important for this activity in *MAT*a cells [25]. We find that Sir2 indirectly supports donor preference from the left half of the RE in *MAT*a cells by silencing *HMLALPHA2* expression, which prevents transcriptional repression by an Mcm1/α2 heterodimer. Similarly, the loss of Sir2 also represses *RDT1* expression and condensin recruitment in the right half of the RE due to *HMLALPHA2* expression. However, Sir2 directly represses *RDT1* through localized histone deacetylation. How the loss of *RDT1* regulation and condensin recruitment changes chromosome III structure in the *sir2*Δ mutant remains unknown, but we propose that the *HMR-MAT*a interaction is a default state, while the *HML-HMR* association has to be actively maintained by condensin and likely additional factors co-localized to this element.

Interestingly, there also appears to be a function relationship between the RE and silencing at the *HML* locus, such that deleting the left half of the RE specifically stabilizes *HML* silencing in *MAT*a cells [36]. The mechanism involved remains unknown, but we hypothesize that eliminating this part of the RE could potentially allow the SIR and condensin complexes bound at the *RDT1* promoter encroach and somehow enhance the heterochromatic structure at *HML*. Under this scenario, the left half of the RE could be insulating *HML* from the chromosomal organizing activity that occurs at the *RDT1* promoter.

### Condensin function in mating-type switching

The *RDT1* promoter was a major condensin binding site identified by ChIP-seq (Fig 1), and given the strong Hi-C interaction between nearby *HML* and the *HMR* locus, we initially hypothesized that condensin at the *RDT1* promoter would crosslink with another condensin site bound on the right arm of chromosome III. ChIP-seq of Smc4-myc did not reveal any strong peaks near *HMR*, but condensin was clearly enriched at *CEN3* (data not shown). Interestingly, the *S. cerevisiae* condensin complex was recently shown to catalyze ATP-dependent unidirectional loop extrusion using an *in vitro* single molecule assay [37]. The mechanism involves direct binding of condensin to DNA, followed by one end of the bound DNA being pulled inward as an extruded loop. Applying this model to the strong binding site at the *RDT1* promoter, this region could act as an anchor bound by condensin, with DNA to the right being rapidly extruded as a loop until pausing at *CEN3*. Extrusion would then continue at a slower rate toward *HMR*, allowing *HML* the time to sample the entire right arm of chromosome III, until clustering with *HMR*. HOMER analysis of the Hi-C data in Fig 5B provides evidence for such a model because there is an ascending gradient of *HML* interaction frequency with sequences extending from the centromere region (bin 2) toward *HMR*, suggesting that *HML* “samples” the right arm of chromosome III. Once brought in contact, *HML* and *HMR* would then remain associated due to their heterochromatic states and shared retention at the nuclear envelope [38] In addition to preventing *HMR* association with *MAT*a, we hypothesize that the looped chromosome III structure makes the chromosome licensed for mating-type switching in response to the HO-induced DSB during G1.

### MATa specific recruitment of Sir2 and condensin to the RE

Condensin, and Sir2 each strongly associated with the *RDT1* promoter exclusively in *MAT*a cells, though it is not clear if they bind at the same time, or are differentially bound throughout the cell cycle. Since DPS2 was required for Sir2 and condensin recruitment, and derepression of *HMLALPHA2* from *HML* also eliminated binding, we hypothesized and then demonstrated (Fig S3) that Mcm1 was a key DNA binding factor involved. Mcm1 is a prototypical MADS box combinatorial transcription factor that derives its regulatory specificity through interactions with other factors, such as Ste12 in the case of *MAT*a haploid-specific gene activation, or α2 when repressing the same target genes in *MAT*α cells [39]. This raises the question of whether Mcm1 directly recruits the SIR and condensin complexes, or perhaps additional factors that work with Mcm1 are involved. At the *RDT1* promoter, specificity for Sir2/condensin recruitment could originate from sequences underlying the condensin/Sir2 peaks. There are no traditional silencer-like sequences for SIR recruitment within the deleted 100bp (coordinates 30702 to 30801), and yeast condensin does not appear to have a consensus DNA binding sequence [40]. Closer inspection of the *RDT1* promoter indicates an A/T rich region with consensus binding sites for the transcription factors Fkh1/2 and Ash1, each of which regulates mating-type switching [11, 41, 42]. Fkh1 and Fkh2 also physically associate with Sir2 at the mitotic cyclin *CLB2* promoter during stress [43]. Ash1 is intriguing because it represses HO transcription in daughter cells [42, 44], raising the possibility of *RDT1* repression in daughter cells. Mcm1 activity in *MAT*a cells could also indirectly establish a chromatin environment that is competent for Sir2/condensin recruitment, rather than direct recruitment through protein-protein interactions. In *MAT*a cells, Mcm1 appears to prevent the strong nucleosome positioning across the RE that occurs in *MAT*α cells [25], and indicative of an actively remodeled chromatin environment. Perhaps condensin is attracted to such regions, which is consistent with the association of condensin with promoters of active genes in mitotic cells, where enrichment was greatest at unwound regions of DNA [45]. Furthermore, nucleosome eviction by transcriptional coactivators was found to assist condensin loading in yeast [46], though the mechanism of loading remains poorly understood. Recruitment of condensin to the *RDT1* promoter LCR therefore provides an outstanding opportunity for dissecting mechanisms of condensin loading and function.

## Methods

### Yeast strains, plasmids, and media

Yeast strains were grown at 30°C in YPD or synthetic complete (SC) medium where indicated. The *SIR2*, or *HST1* open reading frames (ORFs) were deleted with *kanMX4* using one-step PCR-mediated gene replacement. *HML* was deleted and replaced with *LEU2*. A 100bp deletion within the *RDT1* promoter (chrIII coordinates 30701-30800) or DPS2 deletion (chrIII coordinates 30557-30626) was generated using the *delitto perfetto* method [47]. Endogenous *SIR2, BRN1*, or *SMC4* genes were C-terminally tagged with the 13xMyc epitope (13-EQKLISEEDL). Deletion and tagged genes combinations were generated through genetic crosses and tetrad dissection, including Brn1 tagged with a V5-AID tag (template plasmids kindly provided by Vincent Guacci). All genetic manipulations were confirmed by PCR, and expression of tagged proteins confirmed by western blotting. The pGAL-HO-URA3 expression plasmid was kindly provided by Jessica Tyler [27]. Strain genotypes are provided in Supplemental Table S2 and oligonucleotides listed in Table S3.

### ChIP-Seq analysis

Sir2 ChIP-seq was previously described [19]. For other ChIP-seq datasets, log-phase YPD cultures were cross-linked with 1% formaldehyde for 20 min, pelleted, washed with Tris-buffered saline (TBS), and then lysed in 600 µl FA140 lysis buffer with glass beads using a mini-beadbeater (BioSpec Products). Lysates were removed from the beads and sonicated for 60 cycles (30s “on” and 30s “off” per cycle) in a Diagenode Bioruptor. Sonicated lysates were pelleted for 5 min at 14000 rpm in a microcentrifuge and the entire supernatant was transferred to a new microfuge tube and incubated overnight at 4°C with 5 µg of anti-Myc antibody (9E10) and 20 µl of protein G magnetic beads (Millipore). Following IP, the beads were washed once with FA140 buffer, twice with FA500 buffer, and twice with LiCl wash buffer. DNA was eluted from the beads in 1% SDS/TE buffer and cross-links were reversed overnight at 65°C. The chromatin was then purified using a Qiagen PCR purification kit. Libraries were constructed using the Illumina Trueseq ChIP Sample Prep kit and TrueSeq standard protocol with 10ng of initial ChIP or Input DNA. Libraries that passed QC on a Bioanalyzer High Sensitivity DNA Chip (Agilent Technologies) were sequenced on an Illumina Miseq (UVA DNA Sciences Core).

### ChIP-seq computational analysis

Biological duplicate fastq files were concatenated together and reads mapped to the sacCer3 genome using Bowtie with the following options: ---best, --stratum, --nomaqround, and --m10 [48]. The resulting bam files were then converted into bigwig files using BEDTools [49]. As part of the pipeline, chromosome names were changed from the sacCer3 NCBI values to values readable by genomics viewers e.g. “ref|NC_001133|” to “chrI”. The raw and processed datasets used in this study have been deposited in NCBI’s GEO and are accessible through the GEO series accession number GSE92717. Downstream GO analysis was performed as follows. MACS2 was used to call peaks with the following options: --broad, --keep-dup, -tz 150, and -m 3, 1000 [50]. GFP peaks in the WT or *sir2*Δ backgrounds were subtracted from the WT *SMC4-13xMyc* and *sir2*Δ *SMC4-13xMyc* peaks, respectively, using BEDTools “intersect” with the –v option. The resulting normalized peaks were annotated using BEDTools “closest” with the -t all option specified, and in combination with a yeast gene list produced from USCS genome tables. The annotated peaks were then analyzed for GO terms using YeastMine (yeastmine.yeastgenome.org).

### Hi-C analysis

Log-phase cultures were cross-linked with 3% formaldehyde for 20 min and quenched with a 2x volume of 2.5M Glycine. Cell pellets were washed with dH_2_O and stored at −80°C. Thawed cells were resuspended in 5 ml of 1X NEB2 restriction enzyme buffer (New England Biolabs) and poured into a pre-chilled mortar containing liquid N_2_. Nitrogen grinding was performed twice as previously described [51], and the lysates were then diluted to an OD_600_ of 12 in 1x NEB2 buffer. 500 µl of cell lysate was used for each Hi-C library as follows. Lysates were solubilized by the addition of 50 µl 1% SDS and incubation at 65°C for 10 min. 55 µl of 10% TritonX-100 was added to quench the SDS, followed by 10 µl of 10X NEB2 buffer and 15 µl of *Hin*dIII (New England Biolabs, 20 U/µl) to digest at 37°C for 2 hr. An additional 10 µl of *Hin*dIII was added for digestion overnight. The remainder of the protocol was based on previously published work with minor exceptions [52]. In short, digested chromatin ends were filled-in with Klenow fragment (New England Biolabs) and biotinylated dCTP at 37°C for 1 hr, then heat inactivated at 70°C for 10 min. Ligation reactions with T4 DNA ligase were performed at 16°C for a minimum of 6 hr using the entire Hi-C sample diluted into a total volume of 4 ml. Proteinase K was added and cross-links were reversed overnight at 70°C. The ligated chromatin was phenol:chloroform extracted, ethanol precipitated, then resuspended in 500µl dH_2_O and treated with RNAse A for 45 min. Following treatment with T4 DNA polymerase to remove biotinylated DNA ends that were unligated, the samples were concentrated with a Clean and Concentrator spin column (Zymogen, D4013) and sheared to approximately 300bp with a Diagenode Bioruptor. Biotinylated fragments were captured with 30 µl pre-washed Streptavidin Dynabeads (Invitrogen), then used for library preparation. Hi-C sequencing libraries were prepared with reagents from an Illumina Nextera Mate Pair Kit (FC-132-1001) using the standard Illumina protocol of End Repair, A-tailing, Adapter Ligation, and 12 cycles of PCR. PCR products were size selected and purified with AMPure XP beads before sequencing with an Illumina Miseq or Hiseq. Raw and mapped reads deposited at GEO (GSE92717).

### Hi-C computational analysis

Iteratively corrected heatmaps were produced using python scripts from the Mirny lab hiclib library, http://mirnylab.bitbucket.org/hiclib/index.html. Briefly, reads were mapped using the iterative mapping program, which uses Bowtie2 to map reads and iteratively trims unmapped reads to increase the total number of mapped reads. Mapped reads were then parsed into an hdf5 python data dictionary for storage and further analysis. Mapped reads of the same strains were concatenated using the hiclib library’s “Merge” function. Both individual and concatenated mapped reads have been deposited in GEO. Mapped reads were then run through the fragment filtering program using the default parameters as follows: filterRsiteStart(offset=5), filterDuplicates, filterLarge, filterExtreme (cutH=0.005, cutL=0). Raw heat maps were further filtered to remove diagonal reads and iteratively corrected using the 03 heat map processing program. Finally, the iteratively corrected heatmaps were normalized for read count differences by dividing the sum of each row by the sum of the max row for a given plot, which drives all values towards 1 to make individual heatmaps comparable.

Observed/Expected heatmaps were created using HOMER Hi-C analysis software on the BAM file outputs from the iterativemapping program of the hiclib library python package [28]. Tag directories were created using all experimental replicates of a given biological sample and the tbp −1 and illuminaPE options. Homer was also used to score differential chromosome interactions between the WT and mutant Hi-C heatmaps. The resulting list of differential interactions was uploaded into R where the given p-value was adjusted to a qvalue with p.adjust. An FDR cutoff of 0.05 was used to create a histogram of significantly different chromosome interactions in the mutants compared to WT. The histogram was further normalized by dividing the total number of significant differential interactions for a chromosome by total number of interactions called in the WT sample for that chromosome to account for size differences in the chromosomes. Thus, frequency represents the number of interactions that changed out of all possible interactions that could have changed.

### RNA-seq data analysis

RNA-Seq data was acquired from GEO accessions GSE73274 [53] and GSE58319 [54] for the BY4742 (*MAT*α) and BY4741 (*MAT*a) backgrounds, respectively. Reads were then mapped to the sacCer3 genome using Bowtie2 with no further processing of the resulting BAM files visualized in this paper.

### 3C assays

Chromosome Conformation Capture (3C) was performed in a similar manner to Hi-C with a few exceptions due to assay-specific quantification via quantitative real-time PCR rather than sequencing. Most notably, digested DNA ends were not filled in with dCTP-biotin before ligation and an un-crosslinked control library was created for each 3C library. Furthermore, all PCR reactions were normalized for starting DNA concentration using a *PDC1* intergenic region that is not recognized by *Hin*dIII, in addition to PCR of the un-crosslinked control for all tested looping interaction primer pairs.

### Quantitative reverse transcriptase (RT) PCR assay

Total RNA (1 µg) was used for cDNA synthesis with oligo(dT) and Superscript II reverse transcriptase as previously described [55].

### Western blot

Proteins were blotted using standard TCA extraction followed by SDS-PAGE as previously described [19]. Myc-tagged proteins were incubated with an anti-Myc primary antibody 9E10 (Millipore) at a 1:2000 dilution while tubulin was incubated with anti-Tubulin antibody B-5-1-2 (Sigma-Aldrich) at a 1:1500 dilution. The V5-AID tagged Brn1 was detected with anti-V5 antibody (Invitrogen, R96025) at a 1:4000 dilution. Primary antibodies were detected with an anti-mouse secondary antibody conjugated to HRP (Promega) at 1:5000 dilution in 5% fat-free milk. Bands were then visualized with HyGlo (Denville Scientific) capture on autoradiography film (Denville Scientific).

### Mating-type switching assays

For tracking the efficiency of switching, strains were transformed with pGAL-HO-*URA3*, pre-cultured in SC-ura + raffinose (2%) medium overnight, re-inoculated into the same medium (OD_600_=0.05) and then grown into log phase. Galactose (2%) was added to induce HO expression for 45 min. Glucose (2%) was then added and aliquots of the cultures were harvested at indicated time points. Genomic DNA was isolated and 10 ng used for PCR amplification. *MAT*α was detected using primers JS301 and JS302. The *SCR1* gene on chromosome V was used as a loading control (primers JS2665 and JS2666). PCR products were separated on a 1% agarose gel stained with ethidium bromide and then quantified using ImageJ. Donor preference with strains containing *HMR*α-B was performed as previously described [15]. Briefly, MATa was amplified with primers Yalpha105F and MATdist-4R from genomic DNA 90 after switching was completed (90 min), and then digested with BamHI. Ethidium stained bands were quantified using ImageJ. For the conditional V5-AID degron strains, degradation of V5-AID-fused Brn1 protein was induced by addition of 0.5 mM indole-3-acetic acid (Auxin, Sigma # 13750).

## Author contributions

Conceptualization; M.L., R.D.F., and J.S.S.; Methodology, M.L, R.D.F., M.D., and J.S.S.; Software, R.D.F. and S.B.; Strain Creation and Validation, M.L., R.D.F., and M.D., Plasmid Creation and Validation, M.L., and R.D.F.; Formal Analysis, M.L., R.D.F., and S.B.; Data Curation, R.D.F.; Writing-Original Draft, M.L., R.D.F. and J.S.S.; Writing Review & Editing, M.L., R.D.F., M.D., and J.S.S; Supervision, J.S.S. and S.B.; Project Administration, J.S.S.; Funding Acquisition, J.S.S.

## Acknowledgments

We thank Job Dekker, Jon Belton, Maitreya Dunham, Ivan Liachko, Maxim Imakev, and Anton Goloborodko for advice on Hi-C protocols and analysis methods. We also thank James Haber, Andrew Murray, Jasper Rine, Jessica Tyler, Alan Hinnebusch, Marc Gartenberg, Dan Gottschling, and Vincent Guacci for kindly providing yeast strains, plasmids, or antibodies. We thank David Auble and Patrick Grant for reading the manuscript and providing suggestions prior to submission. We declare no conflicts of interest.

## Supporting Figure and Table Captions

**Fig S1. *MAT*a-specific transcription of *RDT1* is repressed by Sir2 and Hst1. (A)** IGV screenshot of compiled raw RNA-seq read data from BY4741 (*MAT*a) and BY4742 (*MAT*α) strains. The top two blue peaks represent Smc4-myc and Sir2-myc ChIP-seq reads. **(B)** Quantitative ChIP assay showing additional SIR complex subunit enrichment at the *RDT1* promoter. **(C)** RT-qPCR showing effects of deleting *SIR2* and/or *HST1* on *RDT1* expression when HML is present or deleted (*p<0.05, **p<.005).

**Fig S2. Deletion of Sir2 or the *RDT1* promoter Sir2/condensin binding site does not affect protein levels of Sir2 or Myc-tagged condensin subunits. (A)** Western blot showing steady state Sir2 protein levels in WT (ML1), *sir2*Δ (ML25), and 100bpΔ (ML275) strains. **(B)** Western blot with anti-Myc detection of Brn1-13xMyc or Sir2 in WT (ML149), *sir2*Δ (ML161), and 100bpΔ (ML322) strains. **(C)** Western blot with anti-Myc detection of Smc4-13xMyc or Sir2 in WT (ML152), *sir2*Δ (ML160), and 100bpΔ version.

**Fig S3. The *RDT1*-proximal Mcm1/a2 binding site (DPS2) is important for Sir2 and condensin recruitment. (A)** Schematic diagram depicting the location of the DPS2 sequence deletion relative to other elements with the RE, with the deleted chromosome III coordinates indicated in red. **(B)** Quantitative ChIP of native Sir2 in WT and *dps2*Δ strains. **(C)** Quantitative ChIP of Brn1-Myc in WT and *dps2*Δ strains. @*RDT1* promoter indicates enrichment at the Sir2/condensin peak (**p<0.005).

**Fig S4. Deleting the Sir2/condensin binding site within the RE (100bpΔ) does not alter Sir2 function at *HML*α. (A)** Quantitative mating assay for WT (ML1) and 100bpΔ (ML275) strains. Quantitative ChIP assay showing Sir2 enrichment at *HML-I* in WT (ML1) and 100bpΔ (ML275) strains. (**p<0.005).

**Fig S5. Auxin inducible degron (AID)-mediated depletion of Brn1 does not derepress *RDT1* or *HML*α. (A)** Western blot time course of auxin induced degradation of Brn1::V5-AID. Time indicates minutes after addition of auxin. **(B)** RT-qPCR of *RDT1* expression following 30 or 60 minutes of Brn1 depletion by auxin. **(C)** RT-qPCR of *HMLALPHA2* expression following 30 or 60 min of Brn1 depletion by auxin.

**Table S1. Genes closest to Sir2-dependent condensin peaks.** This Excel spreadsheet lists the systematic ORF names of all genes that were closest to Sir2-dependent condensin peaks, as chosen using MACS.

**Table S2. Yeast Strains.** List of all Saccharomyces cerevisiae strains used in this study, along with their genotypes and source.

**Table S3. Oligonucleotides.** List of oligodeoxynucleotides used in this study.

