## Supplemental Information for "A Sir2-regulated locus control region in the recombination enhancer of *Saccharomyces cerevisiae* specifies chromosome III structure"

### Supplemental Information (Li et al.)

**Supplemental Table S1. Genes closest to Sir2-dependent condensin peaks.**

**(See Excel file).**

**Supplemental Table S2. Yeast Strains**

| Strains | Genotype | Source |
|---|---|---|
| ML1 | *MAT*a *his3Δ200 leu2Δ1 met15Δ0 trp1Δ63 ura3-167* | [1] |
| ML25B12 | *MATα his3Δ200 leu2Δ1 met15Δ0 trp1Δ63 ura3-167* | This study |
| ML25 | *MAT*a *his3Δ200 leu2Δ1 met15Δ0 trp1Δ63 ura3-167 sir2Δ::kanMX4* | [1] |
| ML26 | *MAT*a *his3Δ200 leu2Δ1 met15Δ0 trp1Δ63 ura3-167 sir3Δ::kanMX4* | [1] |
| ML27 | *MAT*a *his3Δ200 leu2Δ1 met15Δ0 trp1Δ63 ura3-167 sir4Δ::kanMX4* | [1] |
| ML28 | *MATα his3Δ200 leu2Δ1 met15Δ0 trp1Δ63 ura3-167 sir2Δ::kanMX4* | This study |
| ML149 | *MAT*a *his3Δ200 leu2Δ1 met15Δ0 trp1Δ63 ura3-167 BRN1::13xMyc-kanMX4* | [2] |
| ML152 | *MAT*a *his3Δ200 leu2Δ1 met15Δ0 trp1Δ63 ura3-167 SMC4::13xMyc-kanMX4* | [2] |
| ML160 | *MAT*a *his3Δ200 leu2Δ1 met15Δ0 trp1Δ63 ura3-167 SMC4::13xMyc-kanMX4 sir2Δ::natMX4* | [2] |
| ML161 | *MAT*a *his3Δ200 leu2Δ1 met15Δ0 trp1Δ63 ura3-167 BRN1::13xMyc-kanMX4 sir2Δ::natMX4* | [2] |
| ML195 | *MAT*a *his3Δ200 leu2Δ1 met15Δ0 trp1Δ63 ura3-167 sir2Δ::kanMX4* [pRS315] | This study |
| ML196 | *MAT*a *his3Δ200 leu2Δ1 met15Δ0 trp1Δ63 ura3-167 sir2Δ::kanMX4* [pRS315-*SIR2*] | This study |
| ML197 | *MAT*a *his3Δ200 leu2Δ1 met15Δ0 trp1Δ63 ura3-167 sir2Δ::kanMX4* [pRS315-*sir2-H364Y*] | This study |
| ML275 | ML1 deleted for the 100bp Sir2/condensin binding site (100bpΔ) | This study |
| ML279 | ML275 made *sir2Δ::kanMX4* | This study |
| ML286 | ML25B12 deleted for the 100bp Sir2/condensin binding site (100bpΔ) | This study |
| ML322 | ML149 deleted for the 100bp Sir2/condensin binding site (100bpΔ) | This study |

| | | |
|---|---|---|
| ML337 | *MATα his3Δ200 leu2Δ1 met15Δ0 trp1Δ63 ura3-167 BRN1::13xMyc-kanMX4* | This study |
| ML339 | *MATα his3Δ200 leu2Δ1 met15Δ0 trp1Δ63 ura3-167 SMC4::13xMyc-kanMX4* | This study |
| ML341 | *MAT*a *his3Δ200 leu2Δ1 met15Δ0 trp1Δ63 ura3-167 dps2Δ* | This study |
| ML343 | *MAT*a *his3Δ200 leu2Δ1 met15Δ0 trp1Δ63 ura3-167 sir2Δ::kanMX4 hmlΔ::LEU2* | [1] |
| ML344 | *MAT*a *his3Δ200 leu2Δ1 met15Δ0 trp1Δ63 ura3-167 hmlΔ::LEU2* | [1] |
| ML350 | *MAT*a *his3Δ200 leu2Δ1 met15Δ0 trp1Δ63 ura3-167 BRN1::13xMyc-kanMX4 hmlΔ::LEU2* | This study |
| ML351 | *MAT*a *his3Δ200 leu2Δ1 met15Δ0 trp1Δ63 ura3-167 BRN1::13xMyc-kanMX4 sir2Δ::kanMX4 hmlΔ::LEU2* | This study |
| ML419 | *MAT*a *his3Δ200 leu2Δ1 met15Δ0 trp1Δ63 ura3-167 RDT1::13xMyc-kanMX4* | This study |
| ML432 | ML419 deleted for the 100bp Sir2/condensin binding site (100bpΔ) | This study |
| ML433 | *MATα his3Δ200 leu2Δ1 met15Δ0 trp1Δ63 ura3-167 RDT1::13xMyc-kanMX4* | This study |
| ML440 | ML1 [pGAL-HO-*URA3*] | This study |
| ML444 | ML149 [pGAL-HO-*URA3*] | This study |
| ML523 | *MAT*a *his3Δ200 leu2Δ1 met15Δ0 trp1Δ63 ura3-167 BRN1::13xMyc-kanMX4 dps2Δ* | This study |
| ML557 | XW652 *sir2Δ::kanMX4* | This study |
| XW652 | *MAT***a** *ho ade3::GAL::HO HMLαRE HMRα*-B *ura3-52 lys5 leu2-3,112 trp1::hisG* | [3] |
| XW676 | *MAT***a** *ho ade3::GAL::HO HMLαREΔ::URA3 HMRα*-B *ade1 leu2 trp1 ura3-52* | [3] |
| SY742 | XW652 deleted for the 100bp Sir2/condensin binding site (100bpΔ) | This study |
| NBY8 | *MATa ura3-1 leu2-3,112 his3-11 trp1-1 ade2-1 can1-100 bar1Δ lys2Δ* | [4] |
| NBY316 | *MATa ura3-1 leu2-3,112 his3-11 trp1-1 ade2-1 can1-100 bar1Δ lys2Δ ycs4-1* | [4] |
| NBY585 | *MATa ura3-1 leu2-3,112 his3-11 trp1-1 ade2-1 can1-100 bar1Δ lys2Δ hmlΔ::LEU2* | [4] |
| NBY319 | *MATa ura3-1 leu2-3,112 his3-11 trp1-1 ade2-1 can1-100 bar1Δ lys2Δ hmlΔ::LEU2 ycs4-1* | [4] |
| RF15 | *MAT*a *his3Δ200 leu2Δ1 met15Δ0 trp1Δ63 ura3-167*[pRPL25NLS-GFP] | This study |
| RF25 | *MAT*a *his3Δ200 leu2Δ1 met15Δ0 trp1Δ63 ura3-167 sir2Δ::kanMX4*[pRPL25NLS-GFP] | This study |
| MD25 | *MAT*a *leu2Δ1 met15Δ0 trp1Δ63 his3Δ200::pGPD1-Os TIR-* | This study |

| | | |
|---|---|---|
| | *HIS3 BRN1-3V5-AID2:KanMx6* [pGAL-HO-*URA3*] | |
| MD27 | XW652 made *leu2Δ1::pGPD1-Os TIR-LEU2 BRN1::3V5-AID2-KanMX6* | This study |

**Supplemental Table S3. Oligonucleotides**

| Oligo name | Oligo description | DNA sequence |
|---|---|---|
| JS301 | *MAT*α PCR primer-1 | AGTCACATCAAGATCGTTTATGG |
| JS302 | *MAT*α PCR primer-2 | GCACGGAATATGGGACTACTTCG |
| JS467 | *KanMX* 5'-out detection | TACGGGCGACAGTCACATCATG |
| JS1896 | *SPB1* ORF (FW) | CATCGAAGTTAAGGACGACGC |
| JS1897 | *SPB1* ORF (RV) | TCGCGCTTGACATTTAGACG |
| JS1898 | Sir2 binding site 100 bp (FW) | TGTTTGCAAGATGGTGCTTTTT |
| JS1899 | Sir2 binding site 100 bp (RV) | AGGAGCAGAAACGTGGCAAT |
| JS1909 | *HML-I* silencer (ARS302) (FW) | AACATGAAAGCCCGACGTTT |
| JS1910 | *HML-I* silencer (ARS302) (RV) | AATAATCGGGTGAAAAAGAGGATAT |
| JS2127 | *BRN1*_Degron_FW | AGTGAATTATGAGGATCTAGCGACAACACAGGCAGCG TCACGGATCCCCGGGTTAATTAA |
| JS2128 | *BRN1*_Degron_RV | GCACAAAAAAAAAAAAAAAAAAAAAAAAAAAGATCA TCAAGAATTCGAGCTCGTTTAAAC |
| JS2167 | 3C *PDC1* intergenic FWD | GCCGACAGTCTGTTGAATTGG |
| JS2168 | 3C *PDC1* intergenic REV | GAAGCGGACCCAGACTTAAGC |
| JS2342 | Sir2 binding site 100bpΔ pCORE (FW) | GACTTACAAGCACACCTTTGAATTATTTTGTTCTCTAT GTCCTTACCATTAAGTTGATC |
| JS2343 | Sir2 binding site 100bpΔ pCORE (RV) | TATATAGCTATTCATCAATTGAAATATTCATTTTATAAG T GAGCTCGTTTTCGACACTGG |
| JS2444 | pCORE 100bpΔ replacement (FW) | GACTTACAAGCACACCTTTGAATTATTTTGTTCTCTAT GACTTATAAAATGAATATTTCAATTGATGAATAGCTAT ATA |
| JS2445 | pCORE 100bpΔ replacement (RV) | TATATAGCTATTCATCAATTGAAATATTCATTTTATAAG TCATAGAGAACAAAAATAATTCAAAGGTGTGCTTGTA AGTC |
| JS2494 | *RDT1* upstream (-258 bp) (FW) | CGCGTTTAAAGACTTACAAGCAC |
| JS2496 | *RDT1* upstream | TTAAATACATGCTGCAGTTTTCG |

| | | |
|---|---|---|
| | (-44 bp) (RV) | |
| JS2517 | *RDT1* upstream (-19 bp) (FW) | AAAACTGCAGCATGTATTTAATCG |
| JS2518 | *RDT1* downstream (+19 bp) (RV) | TGCTTTCGATTATTTCTGGTTCT |
| JS2574 | Yalpha105F | GCCCACTTCTAAGCTGATTTCAATCTCTCC |
| JS2575 | MATdist-4R | CCTGTTCTTAGCTTGTACCAGAGG |
| JS2583 | *RDT1*-13xMyc_FW | AATTCTATTTGTCCAGCAATCCGGCGCAAAGAAGACTA CCGGATCCCCGGGTTAATTAA |
| JS2584 | *RDT1*-13xMyc_RV | TTTCGATTATTTCTGGTTCTAGAAATTTTTCAATACCCT GAATTCGAGCTCGTTTAAAC |
| JS2585 | *RDT1* detection | TCTATTTGTCCAGCAATCCG |
| JS2656 | 3C *Hin*dIII *HML* | TTCCGAAAACCACGACGAACCAG |
| JS2658 | 3C *Hin*dIII *HMR* | ATGGGTCATTCTAGGTCATTCTAC |
| JS2665 | *SCR1* ORF (33 bp) (FW) | CGTTGAGAATTCTGGCCGAG |
| JS2666 | *SCR1* ORF (468 bp) (RV) | GTAAATCCTGATGGCACCGC |
| JS2669 | KanC3 | CCTCGACATCATCTGCCCAGAT |
| JS2703 | *HO* cut-site 600 bp (FW) | TTGGATCTTAACAAACCGTAAAGGT |
| JS2704 | *HO* cut-site 600 bp (RV) | GGTAACTAGCAAACAAAGGAAAGTCA |
| JS2712 | MATa detection (FW) | TTGCAACAACTTCTTCTCCTCA |
| JS2715 | Alpha2 ORF_FW | TTGGTTTGCAAAGAACATCG |
| JS2716 | Alpha2 ORF_RV | CTTCTTTGCCAGAGGCTCAC |
| JS2777 | R1S/R1L ncRNA (FW) | CGTGCAAACAGTATTCCGGC |
| JS2778 | R1L/R1S ncRNA (RV) | GCGCTGGTTGTTATTGGCAA |

**Supplemental Figure Legends**

**Fig S1.** ***MAT*a-specific transcription of *RDT1* is repressed by Sir2 and Hst1. (A)** IGV screenshot of compiled raw RNA-seq read data from BY4741 (*MAT*a) and BY4742 (*MAT*α) strains. The top two blue peaks represent Smc4-myc and Sir2-myc ChIP-seq reads. **(B)** Quantitative ChIP assay showing additional SIR complex subunit enrichment at the *RDT1* promoter. **(C)** RT-qPCR showing effects of deleting *SIR2* and/or *HST1* on *RDT1* expression when HML is present or deleted (*p<0.05, **p<.005).

**Fig S2. Deletion of Sir2 or the *RDT1* promoter Sir2/condensin binding site does not affect protein levels of Sir2 or Myc-tagged condensin subunits. (A)** Western blot showing steady state Sir2 protein levels in WT (ML1), *sir2*Δ (ML25), and 100bpΔ (ML275) strains. **(B)** Western blot with anti-Myc detection of Brn1-13xMyc or Sir2 in WT (ML149), *sir2*Δ (ML161), and 100bpΔ (ML322) strains. **(C)** Western blot with anti-Myc detection of Smc4-13xMyc or Sir2 in WT (ML152), *sir2*Δ (ML160), and 100bpΔ version.

**Fig S3. The *RDT1*-proximal Mcm1/a2 binding site (DPS2) is important for Sir2 and condensin recruitment. (A)** Schematic diagram depicting the location of the DPS2 sequence deletion relative to other elements with the RE, with the deleted chromosome III coordinates indicated in red. **(B)** Quantitative ChIP of native Sir2 in WT and *dps2*Δ strains. **(C)** Quantitative ChIP of Brn1-Myc in WT and *dps2*Δ strains. @*RDT1* promoter indicates enrichment at the Sir2/condensin peak (**p<0.005).

**Fig S4. Deleting the Sir2/condensin binding site within the RE (100bpΔ) does not alter Sir2 function at *HMLα*. (A)** Quantitative mating assay for WT (ML1) and 100bpΔ (ML275) strains. **(B)** Quantitative ChIP assay showing Sir2 enrichment at *HML-I* in WT (ML1) and 100bpΔ (ML275) strains. (**p<0.005).

**Fig S5. Auxin inducible degron (AID)-mediated depletion of Brn1 does not derepress *RDT1* or *HMLα*. (A)** Western blot time course of auxin induced degradation of Brn1::V5-AID. Time indicates minutes after addition of auxin. **(B)** RT-qPCR of *RDT1* expression following 30 or 60 minutes of Brn1 depletion by auxin. **(C)** RT-qPCR of *HMLALPHA2* expression following 30 or 60 min of Brn1 depletion by auxin.

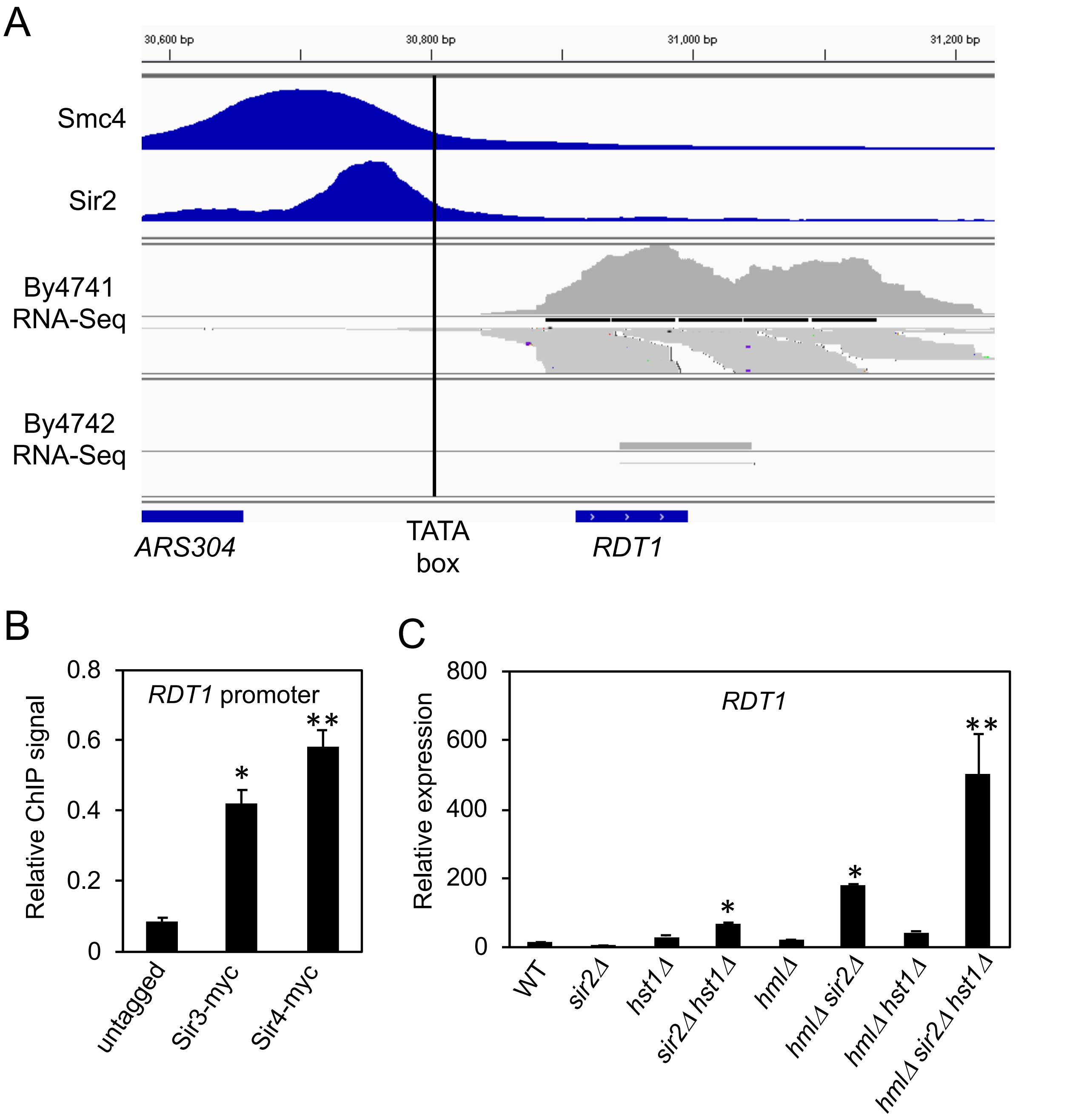
A
30,600 bp
30,800 bp
31,000 bp
31,200 bp
Smc4
Sir2
By4741
RNA-Seq
By4742
RNA-Seq
ARS304
TATA
box
RDT1
B
RDT1 promoter
Relative ChIP signal
0.8
0.6
0.4
0.2
0
*
**
untagged
Sir3-myc
Sir4-myc
C
RDT1
Relative expression
800
600
400
200
0
*
*
**
WT
sir2Δ
hst1Δ
sir2Δ hst1Δ
hmlΔ
hmlΔ sir2Δ
hmlΔ hst1Δ
hmlΔ sir2Δ hst1Δ

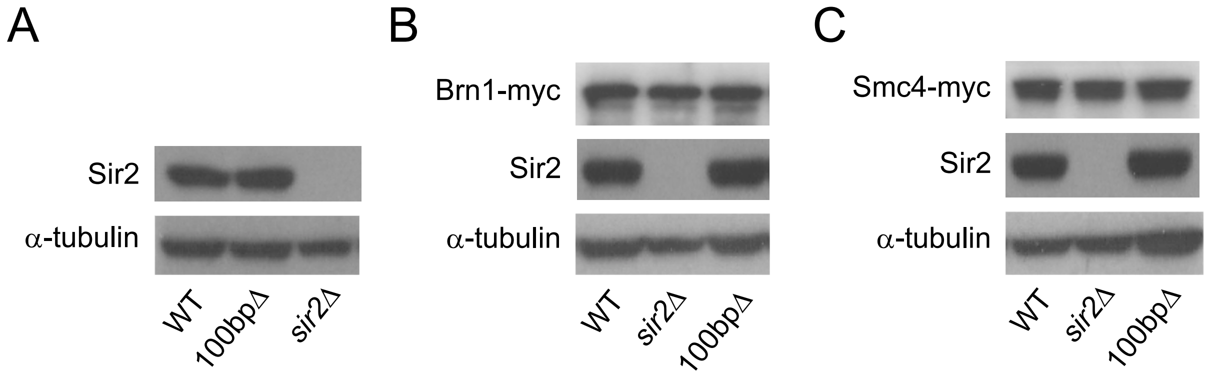
A
Sir2
α-tubulin
WT
100bpΔ
sir2Δ
B
Brn1-myc
Sir2
α-tubulin
WT
sir2Δ
100bpΔ
C
Smc4-myc
Sir2
α-tubulin
WT
sir2Δ
100bpΔ

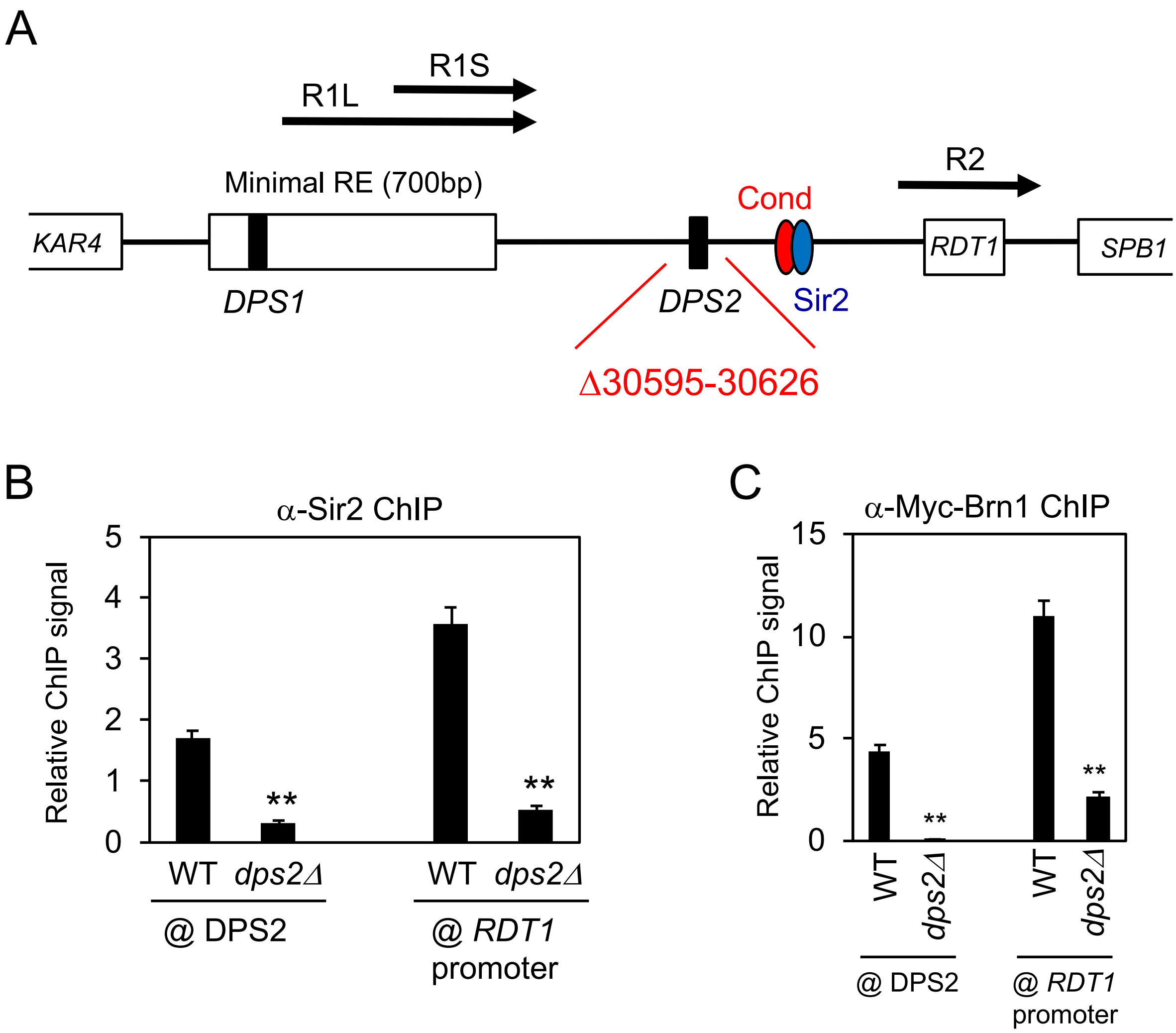
A
R1S
R1L
R2
Minimal RE (700bp)
Cond
KAR4
RDT1
SPB1
DPS1
DPS2
Sir2
Δ30595-30626
B
α-Sir2 ChIP
Relative ChIP signal
5
4
3
2
1
0
**
**
WT dps2Δ
WT dps2Δ
@ DPS2
@ RDT1 promoter
C
α-Myc-Brn1 ChIP
Relative ChIP signal
15
10
5
0
**
**
WT
dps2Δ
WT
dps2Δ
@ DPS2
@ RDT1 promoter

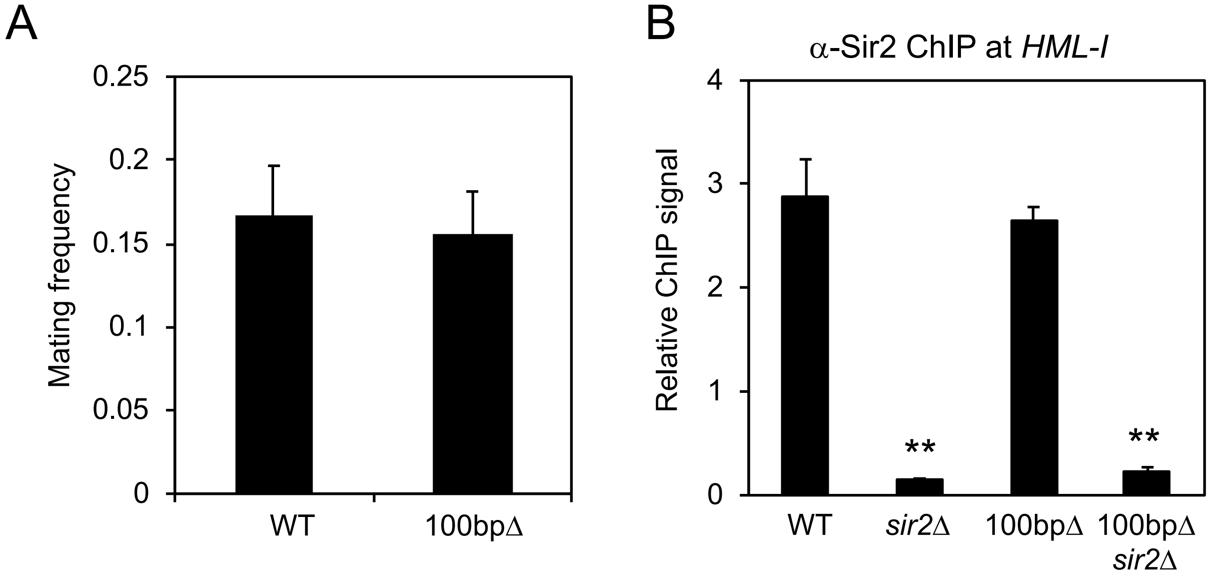
A
0.25
0.2
0.15
0.1
0.05
0
Mating frequency
WT
100bpΔ
B
α-Sir2 ChIP at *HML-I*
4
3
2
1
0
Relative ChIP signal
**
**
WT
*sir2*Δ
100bpΔ
100bpΔ *sir2*Δ

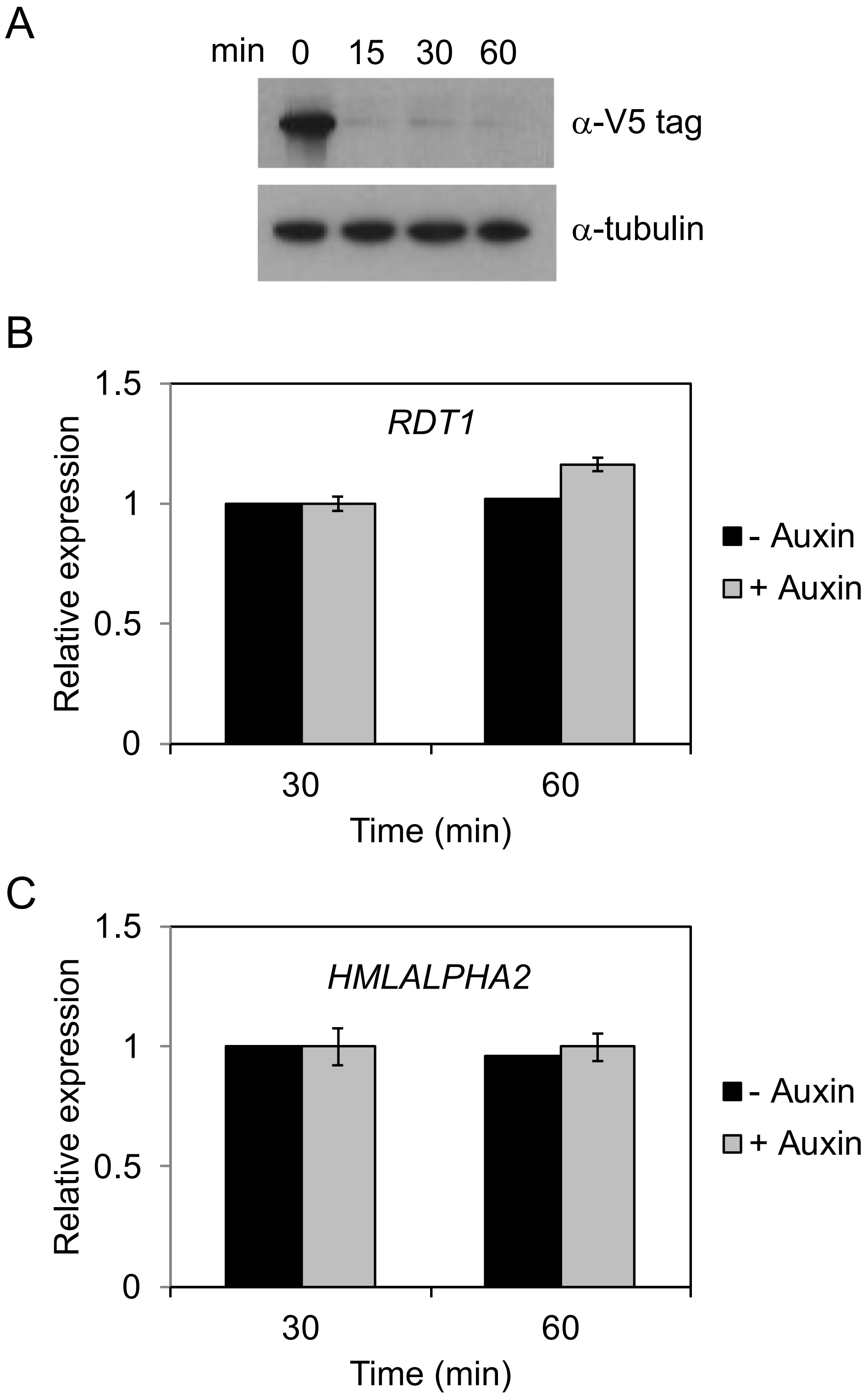
A
min 0 15 30 60
α-V5 tag
α-tubulin
B
RDT1
Relative expression
1.5
1
0.5
0
30
60
Time (min)
- Auxin
+ Auxin
C
HMLALPHA2
Relative expression
1.5
1
0.5
0
30
60
Time (min)
- Auxin
+ Auxin
